# Differential adenosine to inosine RNA editing of SINE B2 non-coding RNAs in mouse unveils a novel type of epi-transcriptome response to amyloid beta neuro-toxicity

**DOI:** 10.64898/2026.01.30.702824

**Authors:** Liam Mitchell, Luke Saville, Travis Haight, Riya Roy, Cody Turner, Yubo Cheng, Igor Kovalchuk, Majid H. Mohajerani, Athanasios Zovoilis

## Abstract

Alzheimer’s disease (AD) is characterized by early molecular responses to amyloid beta neuro-toxicity that remain poorly defined. RNA editing by adenosine-to-inosine (A-to-I) conversion is a major epitranscriptomic mechanism, yet its contribution to non-coding RNA regulation during neurodegeneration is largely unknown. Short Interspersed Nuclear Element (SINE)–derived RNAs, including mouse B2 RNAs, represent the dominant substrates of A-to-I editing and have recently been shown to regulate gene expression through ribozyme-mediated processing. Here, we introduce and validate a repeat-aware bioinformatics framework that enables position-specific quantification of A-to-I editing within highly repetitive SINE RNAs—an analysis that has been challenging using standard genome-based pipelines. Applying this approach, we uncover discrete editing hotspots in B2 RNAs whose modification is selectively increased during the earliest stages of amyloid beta pathology. Elevated B2 RNA editing is consistently observed across hippocampal or neocortical tissue from two independent mouse models of amyloid beta accumulation and in hippocampal neurons exposed to acute amyloid beta toxicity. Functional perturbation of ADAR activity alters both B2 RNA editing levels and the expression of B2 RNA regulated stress-response genes, directly linking RNA editing to SINE-mediated transcriptional control. Independent validation using Nanopore direct RNA sequencing indicates increased RNA modification signals at the same B2 RNA regions identified by short-read Illumina sequencing. Together, our findings establish a previously unrecognized epitranscriptomic response to amyloid beta neurotoxicity mediated by site-specific A-to-I editing of SINE RNAs. This work defines a new analytical paradigm for studying RNA editing in RNAs from repetitive elements and reveals a regulatory axis connecting amyloid beta toxicity, ADAR activity, SINE RNA editing, and stress-responsive gene expression, with implications for conserved mechanisms underlying Alzheimer’s disease and related neurodegenerative disorders.

## INTRODUCTION

Alzheimer’s disease (AD) constitutes one of the most frequent neurodegenerative and aging-associated diseases. A large number of studies have elucidated various cell pathology aspects of AD, with amyloid beta neuro pathology being one of the required features for the diagnosis of AD, and amyloid beta peptides being associated with AD pathogenesis in both human and rodent models (Bloom, 2014; Gonzalez et al., 2018; Ittner et al., 2010; Mehla et al., 2019). However, many of the molecular processes underlying the cellular response to amyloid beta in neural cells remain elusive, which potentially hampers efforts for better molecular diagnostics and therapeutics for this debilitating disease (Bertram & Tanzi, 2008; Hane et al., 2017; Korczyn, 2012; Toledo, Shaw, & Trojanowski, 2013). Among molecular processes that have attracted limited attention to date are potential epi-transcriptome changes associated with amyloid beta pathology.

Currently, across the various domains of life, at least 163 different RNA modifications have been described constituting a significant part of what is known as the epi-transcriptome (Boccaletto et al., 2018). However, many of their functions are not well understood (S. Li & Mason, 2014). A common type of RNA modification in mammalian epi-transcriptomes is RNA editing involving adenosine to inosine conversions (A-to-I editing) (Levanon et al., 2004) (Khermesh et al., 2016; Nakahama & Kawahara, 2020; Neeman, Levanon, Jantsch, & Eisenberg, 2006; Wang, Li, Qi, & Billiar, 2017). A-to-I editing is catalyzed by the ADAR (adenosine deaminases acting on RNA) family of enzymes, which act upon dsRNA structures (Nishikura, 2016; Stefl, Xu, Skrisovska, Emeson, & Allain, 2006; Wagner, Smith, Cooperman, & Nishikura, 1989) and catalyze the conversion of adenosine to inosine by hydrolytic deamination (Bass & Weintraub, 1988; Wagner et al., 1989), with ADAR 1, encoded in mouse by the *Adar* gene, and ADAR2, encoded by the *Adarb1* gene, being the ones responsible for the vast majority of A-to-I editing (ADAR3 encoded by Adarb2 lacks such activity) (Hundley & Bass, 2010; Kapoor et al., 2020; Lehmann & Bass, 2000). These A-to-I edits have been shown to influence directly and indirectly the stability of RNA (Morita et al., 2013; Nishikura, 2016; Solomon et al., 2017) and act as signals that distinguish endogenous RNAs from viral ones and protect them from cellular defense mechanisms and destabilization (Liddicoat et al., 2015; Samuel, 2012; Tomaselli, Galeano, Locatelli, & Gallo, 2015). Additionally, deficiencies in A-to-I RNA editing have been associated with diseases, including neurodegenerative diseases (Nishikura, 2016). In particular, RNA editing in protein coding genes has been shown to alter codon usage of genes associated with synaptic function (Singh, 2012), Alzheimer’s disease (Annese et al., 2018; Khermesh et al., 2016), prion diseases (Kanata et al., 2019) and other neurodegenerative diseases (Krestel & Meier, 2018; Larsen, Hunnicutt, Larsen, Yoder, & Saunders, 2018). However, beyond recoding, little is known about the role of A-to-I editing of non-coding RNAs and particularly SINE RNAs in amyloid pathology, despite SINE RNAs constituting the primary constituents of A-to-I editing in mammalian transcriptomes (Kim et al., 2004; Larsen et al., 2018; Neeman et al., 2006; Paz-Yaacov et al., 2010; Porath, Knisbacher, Eisenberg, & Levanon, 2017).

SINE RNAs are non-coding RNAs generated by repetitive genomic elements called Short Interspersed Nuclear Elements (SINEs). SINEs include B2 elements in mice and Alu elements in humans, which are among the most frequent repeats in the respective genomes and are transcribed either independently by RNA Polymerase III or as part of RNA Polymerase II transcripts in which they are embedded. SINE RNAs constitute one of the primary targets of A-to-I editing (Chen & Carmichael, 2009; Huang et al., 2018; Levanon et al., 2004; Neeman et al., 2006; Porath et al., 2017), which is a process that has been described as essential in marking the RNA as endogenous and, thus, not eliciting an endogenous interferon response as viral RNAs will (Wang et al., 2017). Despite being considered initially as genomic parasites due to their transposon origin (Karijolich, Zhao, Alla, & Glaunsinger, 2017), multiple recent studies have revealed important functions for SINE genomic elements ranging from generation of new regulatory genomic elements to cryptic splicing sites and RNA circularization (Jeck et al., 2013; Kaaij, Mohn, van der Weide, de Wit, & Buhler, 2019; Lin et al., 2008; X. O. Zhang, Gingeras, & Weng, 2019). The same applies to SINE RNAs produced by these genomic elements, which have initially been considered as transcriptional noise generated by the activation of these transposons during cellular stress. However, earlier work from the Goodrich and Kugel labs demonstrated the ability of SINE RNAs to bind and suppress RNA Polymerase II and, thus, marked SINE RNAs as key regulators of gene expression (Espinoza, Goodrich, & Kugel, 2007; Mariner et al., 2008; Ponicsan, Kugel, & Goodrich, 2010, 2015; Yakovchuk, Goodrich, & Kugel, 2009). This activity mediates the transcriptome-wide suppression of housekeeping genes in response to stress, thereby facilitating the redistribution of resources towards pro-survival pathways (Mariner et al., 2008).

Subsequently, we revealed that SINE RNAs regulate the activation of gene expression during cellular response to stress. By occupying stress response genes (B2 SRGs) and suppressing RNA Polymerase II in the pre-stimulus state, B2 RNAs keep these genes poised for fast activation during stress (Zovoilis, Cifuentes-Rojas, Chu, Hernandez, & Lee, 2016). Upon application of a stress stimulus, B2 RNAs, which are self-cleaving RNAs (Hernandez et al., 2020), interact with proteins such as Ezh2 and Hsf1 (Cheng, Saville, Gollen, Isaac, Mehla, et al., 2020; Zovoilis et al., 2016), which accelerate their processing, leading to the release of RNA Polymerase II suppression and the activation of pro-survival genes. As a result, SINE RNAs in mouse act as transcriptional switches during the response to stress stimuli by binding RNA Polymerase II at several stress response genes in the pro-cellular stress state and suppressing their transcription.

Interestingly, in a previous study we found that the above B2 RNA-dependent mode of gene regulation is impaired in hippocampal tissues during amyloid beta pathology. In particular, amyloid toxicity leads to increased destabilization of B2 RNAs in the absence of a stimulus, and, thus, to persistent hyperactivation of B2 regulated stress response genes (B2 SRGs), which in turn are associated with activation of pro-apoptotic pathways (Cheng, Saville, Gollen, Isaac, Mehla, et al., 2020). Given the role of A-to-I editing in regulating RNA stability and that A-to-I editing has been associated with SINE RNAs, we decided to investigate how exactly A-to-I editing is distributed across B2 RNAs, how it is modified in response to amyloid beta pathology and whether it is associated with transcriptome changes in B2 SRGs. However, investigating the potential role of B2 RNA editing in this biological context first requires an analytic framework capable of accurately capturing A-to-I events within these transcripts. The repetitive nature of B2 RNAs and the substantial sequence heterogeneity among individual SINE B2 copies pose major challenges for standard RNA editing analysis pipelines, which were primarily developed for uniquely mappable genomic regions. These limitations hinder reliable quantification of editing at specific base positions of B2 RNAs. Therefore, a second key objective of the present study is to establish a computational strategy tailored to repetitive SINE RNAs that enables robust detection of position-specific A-to-I editing events in B2 RNAs, prior to further examination of these RNAs in the context of pathology.

## RESULTS

### A method for estimating relative A-to-I editing rate across SINE B2 RNAs

For the estimation of A-to-I editing we used Illumina-based next-generation sequencing as the most widely used approach. Using RNA sequencing (RNA-seq), we can interpret A-to-I editing as an A-to-G substitution that occurs during cDNA synthesis during library construction. Data analysis typically involves mapping sequenced reads against the genome and identifying A-to-G sequence conversions relative to the reference sequence that could correspond to potential A-to-I events. The editing rate is then calculated at each genomic position for each transcript. This approach, using, among others tools such as REDI tools (Picardi & Pesole, 2013), has been previously applied to map the exact editing locations in various protein-coding genes and non-repetitive non-coding sequences and in case of RNAs transcribed from repetitive elements to estimate a “bulk” editing rate for entire repeat families (Picardi, D’Erchia, Lo Giudice, & Pesole, 2017). However, in the case of RNAs from repetitive elements such as SINE B2 RNAs, due to their high sequence variability and repetitive nature, such an estimation of their A-to-I editing rate is complicated by a number of parameters. Firstly, SINE B2 RNA sequences are derived from thousands of different loci. The majority of them share identical sequences with other B2 elements located at multiple distinct loci. Thus, mapping of a sequenced read against the genome would often assign a read to multiple locations. To address this problem, many current approaches either filter out reads that are not uniquely mapped to the genome or randomly select only one of the multiple genomics locations that best map the read. However, this approach results in either discarding a large number of potential SINE-mapping reads or misassigning them to the wrong genomic position. The latter could result in SNPs at a specific SINE locus being interpreted falsely as an editing event because B2 elements, despite sharing high sequence similarity with the consensus sequence, also harbor multiple SNPs, small insertions or deletions relative to this sequence.

To circumvent the above problems, we modified the above standard A-to-I analysis approach by adjusting the mapping step and introducing an additional analysis step. An overview of this approach is presented in Fig.1 and described in detail in the methods section. In brief, to account for the obstacles that multiple genomic mapping poses, instead of mapping reads against the mouse reference genome, we have mapped them against a unique “REPome” of SINE B2 sequences. This includes all B2 repetitive elements of a certain family identified by Repeat Masker for assembly version mm10, with the same sequences mapping to multiple loci collapsed into one unique sequence each time. This results in only one sequence with a unique ID regardless of the number of its instances in the genome. In essence, we treat all RNAs generated by multiple different sets of genomic elements sharing the same sequence as a single pool of RNAs studied together. Subsequently, to account for diversity among the different members of the B2 REPome with respect to the B2 consensus sequence, we have implemented an RNA sequence population-based approach. In this approach, instead of calculating the RNA editing for every RNA molecule that is member of the B2 REPome separately, we calculate proportions of base compositions across the entire population of B2 RNAs. This results in the construction of a reference distribution and two sample-specific distributions of base compositions across the length of a B2 RNA metagene model representing all B2 RNAs in the REPome, aligned in with regard to their start position. The reference distribution is the distribution calculated by the reference sequences present in the REPome, independent of the sequenced sample (Fig.1, left plot, annotated as REFERENCE). The second one is the expected distribution if no A-to-I editing occurs, calculated sample-wise from a subset of the first distribution based on the read coverage at each position for each sample (Fig.1, middle plot, annotated as EXPECTED). This distribution is sample-dependent. The third distribution is the actual distribution observed in the reads from each sample (Fig.1, right plot, annotated as OBSERVED) and is also sample-dependent. Based on the last distribution, a ratio of guanosines to the sum of adenosines and guanosines, which represent potential A-to-I events, is calculated for every position and sample. This rate is weighted to the number of expected adenosines at that position to narrow the identification of A-to-I events in positions with frequently occurring adenosines (Fig.1).

**Figure 1.**
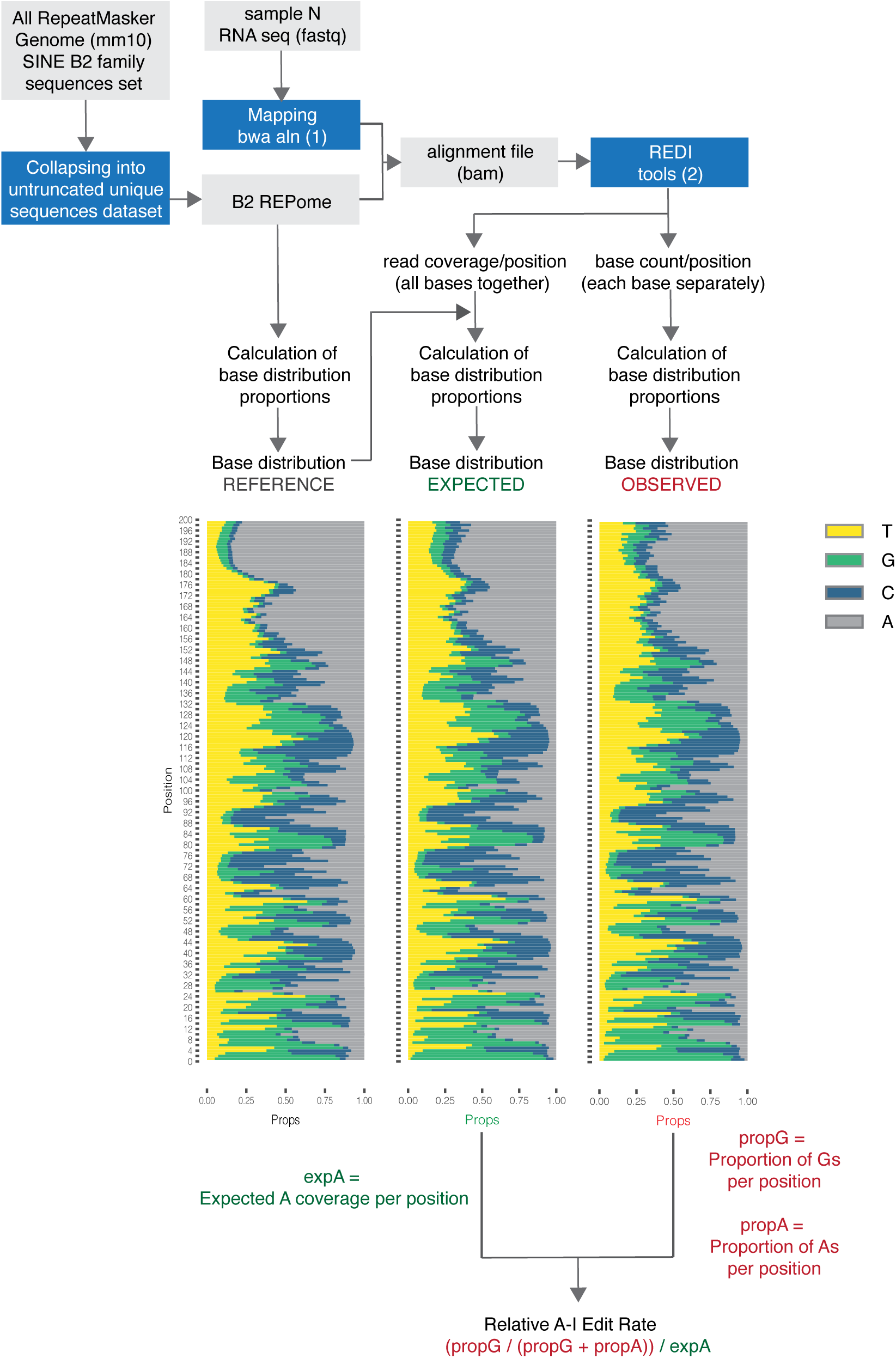
Workflow describing the method for estimating relative A-to-I edit rate across SINE B2 RNAs. Arrows depict the flow of analysis. Plots depict the distribution of base proportions (for A,T,C,G) across B2 RNAs based on i) sequences in the reference (Repeat Masker, mm10) (REFERENCE - left plot), ii) read coverage in each sample with regard to the reference (EXPECTED – middle plot) and iii) actual base coverage observed in each sample (OBSERVED – right plot). Y axis of each plot represents the position in a B2 metagene combining all unique B2 RNA sequences (REPome) aligned at the start site of their consensus sequence with numbers representing the distance from the sequence start (+1). The way the editing rate is calculated is shown in the displayed formula, combining expected and observed base proportions per position and sample. Further details on the calculations of base distributions and of the relative edit rate are provided in Methods. Props: cumulative proportions of four bases (cumulative proportion of all bases added together = 1). (1) as described in (H. Li & Durbin, 2009), (2) as described in (Picardi & Pesole, 2013).

### Adar/Adarb1 knockdown confirms the ability of our customized A-to-I editing analysis to effectively identify A-to-I editing across B2 RNAs

To assess the performance of our customized pipeline in detecting A-to-I editing in SINE B2 RNAs, we performed, in the hippocampal cell line HT-22, siRNA-mediated knockdown of the transcripts of the two main A-to-I editing enzymes, *Adar* for ADAR1 *and Adar1b1* for ADAR2 with their combined knockdown referred as “AntiAdar” throughout the text. The above cell line has previously been used by us and others as a model of amyloid beta induced toxicity in hippocampal-derived neurons (Cheng, Saville, Gollen, Isaac, Belay, et al., 2020; L. Zhang et al., 2018). Cells treated with a scramble siRNA (Scramble) served as a control. The experimental design is outlined in Figure 2A.

**Figure 2.**
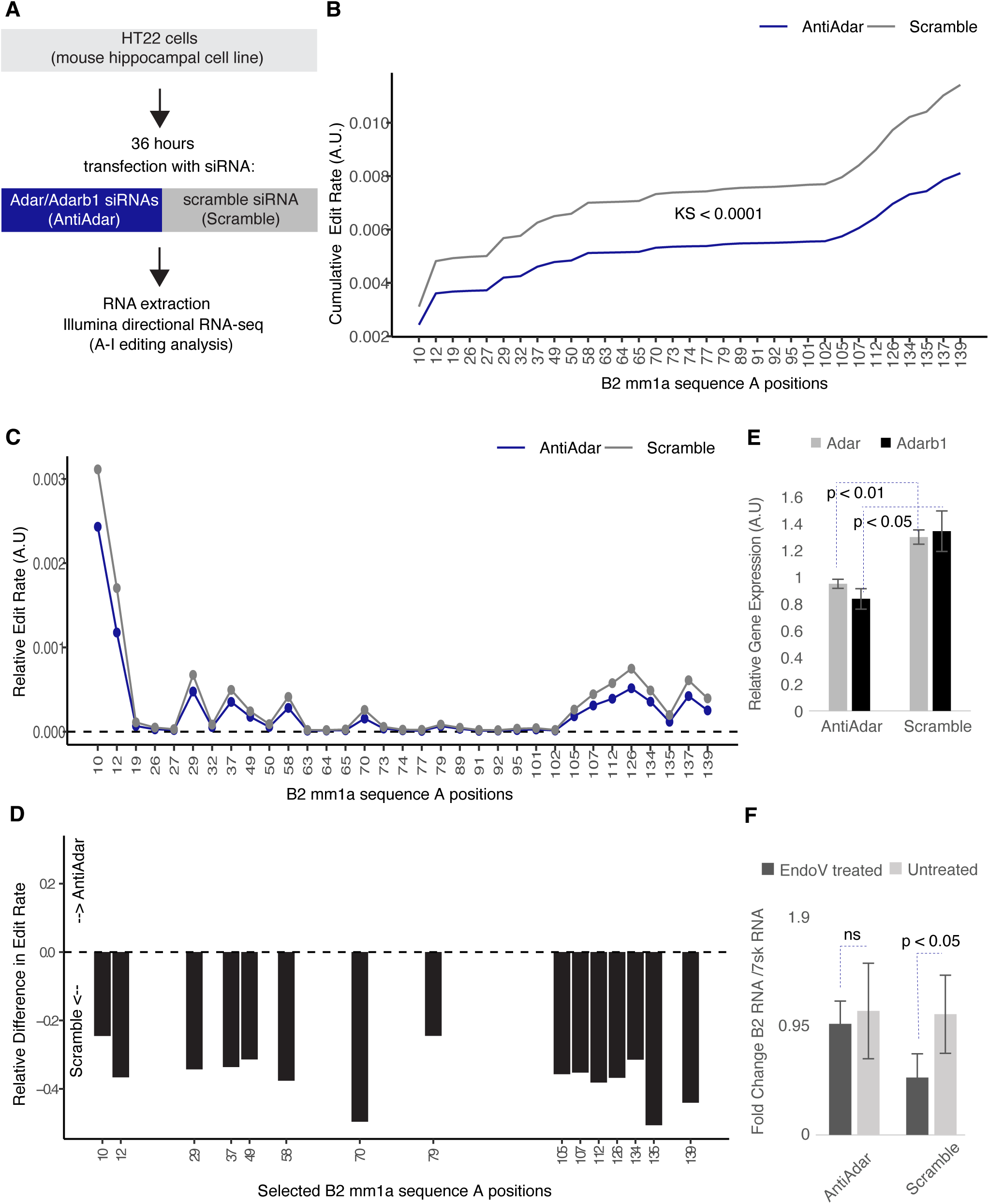
Knockdown of *Adar* and *Adarb1* reduces A-to-I editing of B2 RNAs in hippocampal cells. **(A)** Experimental design for siRNA-mediated knockdown of *Adar* and *Adarb1* (AntiAdar) or scramble control in HT-22 cells (Scramble). Four biological replicates were used per group. **(B)** Cumulative edit rate distributions across B2 RNAs (mm1a) in AntiAdar and Scramble groups (mean per group per position, Kolmogorov–Smirnov test value shown). The X axis shows one value per Adenosine position in the B2 mm1a consensus sequence (A.U.: arbitrary units). **(C)** Position-specific mean relative edit rate across all consensus A positions in the B2 mm1a sequence (A.U.: arbitrary units). **(D)** Relative differences in edit rate (AntiAdar – Scramble) at positions with relative edit rate > 0.0001 (see panel C); negative values denote higher editing in Scramble controls. X axis has one value per Adenosine position in the B2 mm1a consensus sequence. **(E)** Succesful *Adar and Adarb1* KD under the transfection conditions used in this study. qPCR quantification of *Adar and Adarb1* mRNA levels (normalized to *Hprt* expression) in an independent set of experiments.Error bars denote standard deviation (n=3, one-tailed Student’s *t* test). **(F)** Endonuclease V (EndoV) assay showing fold-change differences in B2 RNA levels in Scramble but not AntiAdar samples, measured using RT-qPCR and normalized to *7SK.* Error bars denote standard deviation (n=4 per group, one-tailed Student’s *t* test, ns:non-significant).

Then, using our customized pipeline we calculated the relative A-to-I edit rate at B2 RNA positions for each sample, and the average rate within each group (Scramble, AntiAdar). In this study, we focused specifically on members of the mm1a B2 RNA subfamily, as mm1a—and the closely related to it mm1t subfamily—represent the subfamilies that most faithfully correspond to the canonical B2 RNA consensus previously used in the literature to study B2 RNAs suppressive effect on RNA Polymerase II (Espinoza et al., 2007). Subsequently, we visualized the cumulative distributions of A-to-I editing across all those B2 RNA positions containing an adenosine -the potential substrate for ADAR1/2 activity- and confirmed that AntiAdar treated samples displayed significantly lower relative edit rates compared to scramble siRNA controls (Scramble) (Figure 2B). Complementing this cumulative view, Figure 2C depicts relative edit rates at each individual adenosine position, highlighting discrete peaks corresponding to specific editing sites. Positions of these peaks (corresponding to a relative edit rate >10^−4^) are shown in Figure 2D. Other positions outside these hotspots did not exhibit significant relative edit rates (< 10^−4^ ) and were not analyzed further. The amplitude of the peaks at the identified hotspots was markedly reduced following AntiAdar treatment depicting a relative difference between AntiAdar and Scramble between -0.2 and -0.5, confirming that the identified A-to-I conversions at these positions depend on ADAR1/2 activity (Fig. 2D).

Knockdown efficiency was validated by RT-qPCR, confirming downregulation of both *Adar* and *Adarb1* transcripts (Figure 2E). To further substantiate these results, we performed an assay using Endonuclease V (EndoV), which specifically cleaves inosine-containing RNAs (Morita et al., 2013). The processed products were then assessed by RT-qPCR against B2 RNAs. In Scramble controls, EndoV treatment reduced full-length B2 RNA levels (normalized to 7SK) whereas this reduction was abolished in AntiAdar treated cells, consistent with the depletion of inosine-bearing transcripts (Figure 2F). Overall, the successful knockdown of *Adar* and *Adarb1* transcripts, which was also confirmed through RNA-seq (Suppl. Figure 1), was highly correlated with changes in relative edit rates of B2 RNA transcripts (Suppl. Figure 1).

The positional distribution of editing events identified by our analysis reveals non-uniformity of A-to-I editing along the B2 RNA as editing was concentrated at specific hotspots, that are often clustered together. Interestingly, several of the observed peaks/peak clusters coincide with B2 RNA regions predicted in previous studies to form double-stranded RNA structures (Espinoza et al., 2007; Podbevsek et al., 2018; Sharma et al., 2024), particularly within the first ∼75 nucleotides and in double-strand domains near the 3′ end.

Finally, we compared the efficiency of our repeat-based, customized pipeline for identifying A-to-I editing across B2 RNAs with that of the standard genome-based approach. As shown in Supplementary Figure 2A, standard genome-based alignment (used as a first step for capturing editing events) failed to capture many reads that our B2 REPome (B2ome)-based alignment method was able to capture. Supplementary Figure 2B illustrates how our customized approach accurately resolves position-specific differences in editing between treated and control cells compared to that of the standard genome-based method (Suppl. Figure 2C) which produces a potentially artificial, homogeneous editing pattern (depicted as an almost straight line in the cumulative distribution), lacking the distinct positional differences that would be expected from ADAR-dependent editing activity. It also fails to accurately quantify the editing differences between treated and untreated samples. This loss of positional specificity and overall sensitivity stems from the standard method’s inability to distinguish true editing from sequence variation among the numerous B2 repeat copies.

These results underscore the improved accuracy of our customized method for quantifying A-to-I editing in repetitive elements such as B2 RNAs.

### B2 RNA editing is increased during the first stages of amyloid beta pathology in the mouse hippocampus

Using our customized SINE A-to-I editing analysis, we then investigated whether and how B2 RNA editing levels change in response to amyloid beta pathology. For this purpose, we used the same transcriptome data set used in our previous study for establishing the relationship between B2 RNA processing, target gene hyperactivation and amyloid beta pathology in the mouse hippocampus (Cheng, Saville, Gollen, Isaac, Mehla, et al., 2020). This data set comprises directional Illumina RNA-seq data from the hippocampus of a transgenic mouse model of amyloid beta pathology APP^NL-G-F^ (APP) (Saito et al., 2014), compared with the corresponding wild-type control (C57BL/6J) (WT). This mouse model has been extensively described by us and others with regard to the progression of learning impairment over time and tested for amyloid beta load, RNA expression profiles and B2 RNA processing ratio (Cheng, Saville, Gollen, Isaac, Mehla, et al., 2020; Mehla et al., 2019). The studied APP mice (and the respective WT ones) represented three different stages in the amyloid beta pathogenesis: i) 3 months old (3m, pre-symptomatic stage) ii) 6 months old (6m, start of symptoms - active neurodegeneration phase) and iii) 12 months old (12m, terminal stage - extensive brain atrophy) (Fig.3A).

**Figure 3.**
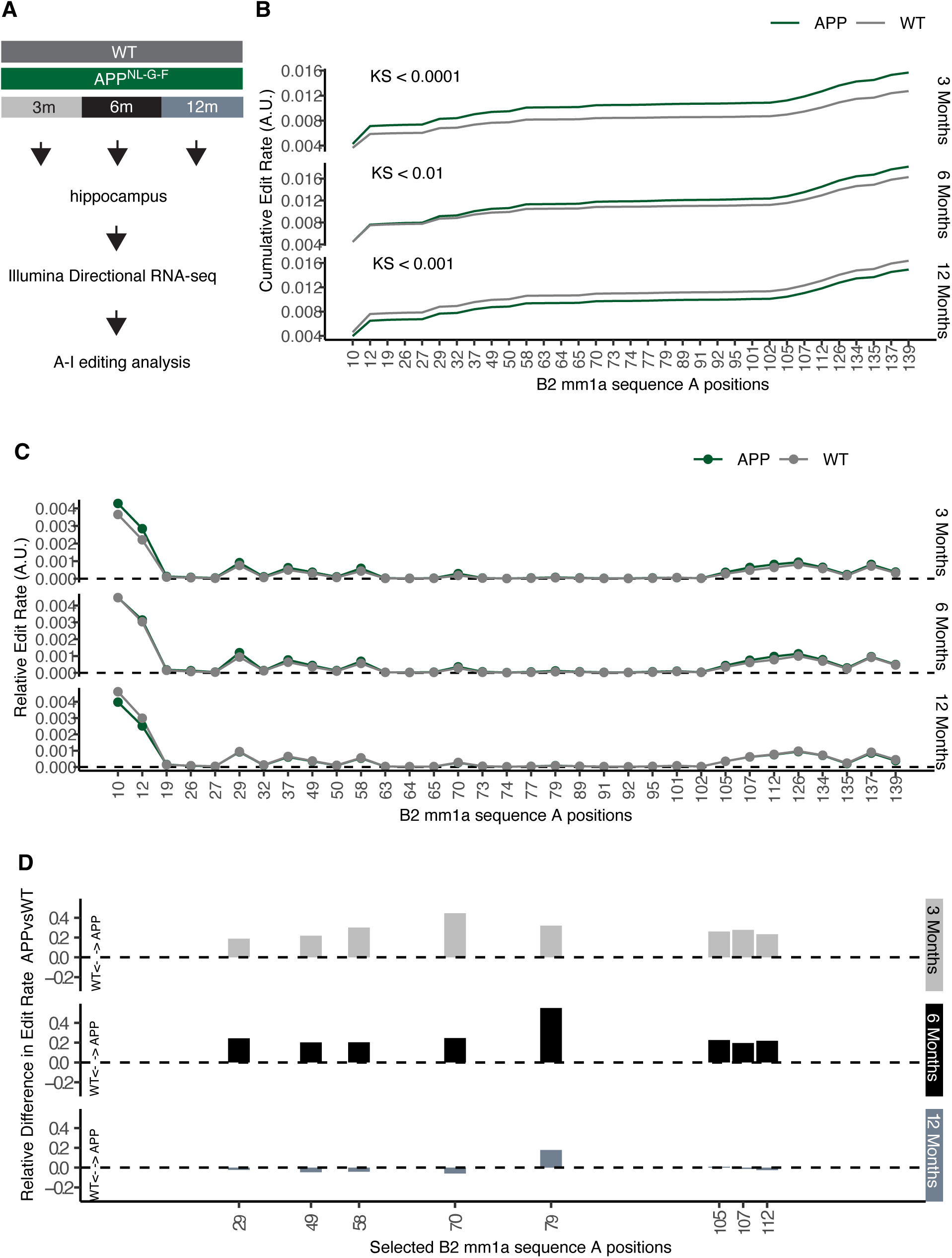
A-to-I editing increases at specific B2 RNA positions during early amyloid beta pathology. **(A)** Experimental overview of hippocampal RNA-seq datasets from APP^NL-G-F^ transgenic (APP) and wild-type (WT) mice at 3, 6, and 12 months of age from (Cheng, Saville, Gollen, Isaac, Mehla, et al., 2020). m denotes the age of mice in months (n=3 per treatment at each age group, except for 3-month WT mice, which were n=2). **(B)** Cumulative distributions of editi rates across B2 RNAs (mm1a) at each time point (Kolmogorov–Smirnov test values shown) (A.U.: arbitrary units). **(C)** Position-specific mean relative edit rates in APP and WT hippocampi at each age (A.U.: arbitrary units). **(D)** Relative differences (APP – WT) in edit rates at consensus A positions of Figure 2D that differ significantly between APP and WT (Suppl. Fig.3). Positive values indicate higher editing in APP.

As in the *Adar1* and *Adarb1* knockdown experiments (AntiAdar), we calculated the relative A-to-I edit rate at each B2 RNA position for each sample, and then the relative edit rate per position among samples within the same age and genotype group. Subsequently, in order to compare the A-to-I edit rate of all positions between APP and WT mice for each age group, we generated cumulative distribution plots of A-to-I editing across all B2 RNA positions containing adenosines (Fig.3B). The resulting distributions suggested an increase in the relative edit rate in APP mice at 3m and 6m compared to WT mice of the same age (KS < 0.0001 and KS < 0.01, respectively), but this increase was eliminated and reversed in 12 months old mice. As shown in Figure 3C, the identified B2 RNA positions with significant relative edit rates (> 10^−4^) correspond to the same positions identified in our AntiAdar experiments in Figure 2C. However, not all of these positions were found to differ significantly between APP and WT mice at 3m and 6m. As shown in Supplementary Figure 3, eight positions showed significant differences, and the relative change in editing rate of these positions is plotted in Figure 3D. This analysis confirmed an increase in A-to-I edit rate in eight positions in 3m and 6m APP mice (Suppl. Fig.3, Fig.3D) while differences in edit rate in the rest of the positions did not reach the statistical significance threshold in either both or one direction comparison in 3m or 6m old mice.

To validate our findings in an independent amyloid beta pathology system and a dataset not generated by our laboratory, we applied our customized SINE A-to-I editing analysis to publicly available brain transcriptome data from an additional mouse model of this pathology (APP^B6.APPP/PSI^). This model exhibits progressive amyloid beta accumulation similar to that observed in the APP^NL-G-F^ strain, but with distinct kinetics and genetic background and a predominant neuroinflammatory phenotype (Chintapaludi et al., 2020). Available Illumina RNA-seq data from whole brain tissue included both transgenic and wild-type control animals up to 6 months old (Suppl. Fig. 4A). As with the analysis of the APP^NL-G-F^ strain, we calculated the relative A-to-I edit rate at each B2 RNA position for every sample and averaged the values within each age and genotype group. Cumulative distributions of A-to-I editing across all adenosine-containing positions (Suppl. Fig. 4B) revealed a similar pattern to that observed in our mouse hippocampus dataset: editing levels were increased in brains of transgenic mice of amyloid beta pathology of the same age (KS < 0.0001). The positions with significant relative edit rates (>10⁻⁴) (Suppl. Fig. 4C) overlapped with those detected in the APP^NL-G-F^ model (Suppl. Fig. 4D) with differences in relative edit rate between APP and WT mice of the eight positions identified in the APP^NL-G-F^ model observed also in the APP^B6.APPP/PSI^ model (Suppl. Fig. 5). The replication of these results in an external dataset demonstrates that the observed editing pattern is not model-specific or dataset-dependent but instead represents a consistent molecular feature of amyloid beta-associated neurodegeneration.

Overall, this data shows that, during the first stages of amyloid beta pathology in mouse hippocampus and whole brain tissue, A-to-I editing at specific conserved positions in B2 RNAs is increased.

### Estimation of RNA modification rates using Nanopore direct RNA sequencing

To evaluate, using an independent method, the level of Adenosine modifications estimated by our customized Illumina-based pipeline, we applied Oxford Nanopore Technologies (ONT) direct RNA sequencing, which enables direct measurement of RNA modifications from raw direct RNA-seq signal data. Using a previously described approach for signal-level analysis of non-standard (modified) bases in this type of data (Pratanwanich et al., 2021), we examined hippocampal RNA samples from both our AntiAdar and scramble siRNA-treated cells (Scramble), as well as from the APP^NL-G-F^ (APP) and wild-type (WT) six-month-old mice used in this study.

To ensure comparability with our Illumina editing rate analysis, Nanopore reads were aligned to a unique B2-ome reference, in which identical B2 sequences across genomic loci were again collapsed into a single representative sequence. Raw Nanopore current signals were first aggregated into 5-mer (k-mer) bins using Nanopolish, and subsequently analyzed with Xpore, a statistical model that estimates modification rates at each k-mer position. Only k-mers containing adenosine residues (the potential substrates for A-to-I editing) and with modification rate values below 0.99 were retained, as higher extreme rates could represent artifacts of the model. Additionally, to obtain the exact electric current signal distributions of modified versus non-modified Adenosines in these samples, Nanopore reads aligned to B2 loci were extracted and the resulting reads within the B2-relative genomics coordinate system were used to generate Nanopolish processed signal plots.

Consistent with our Illumina-based findings, the Xpore-computed modification rate distributions (Fig. 4A) and cumulative modification rate plots (Fig. 4B) showed that the frequency of non-standard A base signals was higher in scramble-treated cells (Scramble) than after ADAR activity knockdown (AntiAdar) (KS=0.001), and higher in 6-month-old APP mice than in age-matched WT controls (KS=0.002). This trend mirrors the increased A-to-I editing rates observed in our Illumina analysis. At a position-specific level, analysis of the raw Nanopore signal distribution across the eight positions previously identified as significantly edited revealed clear differences in the signal profiles between experimental groups (Fig. 4C) in all but one of these positions (position 112)

**Figure 4.**
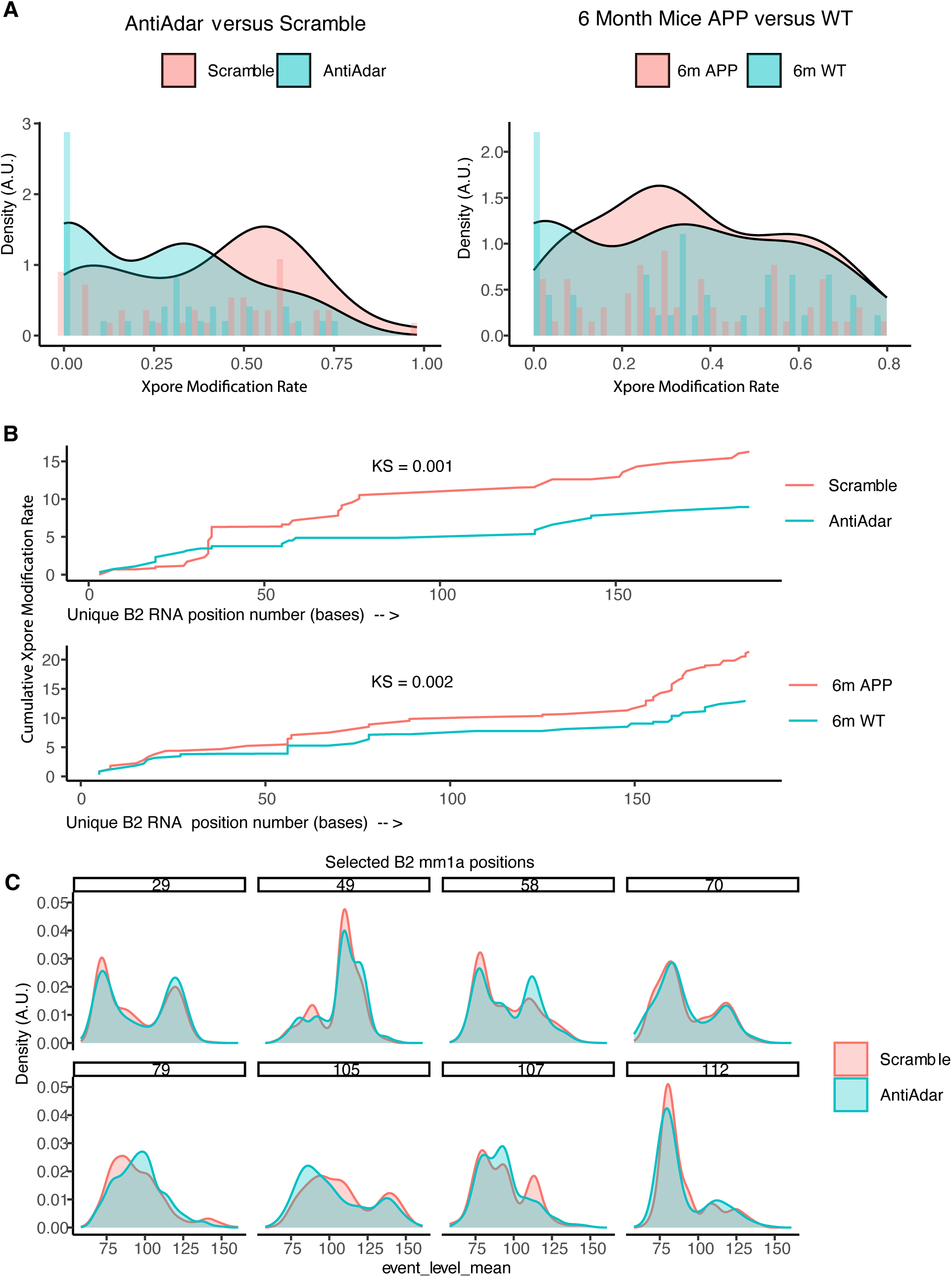
Nanopore direct RNA-seq signal analysis confirms observed RNA modifications in hippocampal cells and mouse hippocampi. (A) Distributions of Xpore-inferred modification rates across B2 RNA positions in AntiAdar versus Scramble HT-22 cells (RNA from four replicates per group pooled per run) and in 6-month-old APP versus WT hippocampi (n=3 per sample group). Samples are concatenated group-wise. X-axis: modification rate bins (overall range 0 [no editing] – 1 [100% editing]); Y axis shows density for the curves and histograms respectively, with histograms scaled down (¼ height) to better display the plot. Density is a smoothed measure of the number of points within a region, and so the points when one sample group rises above another indicates a higher proportion of genomic sites with modification rates within that range. (B) Cumulative modification-rate plots showing aggregate differences in editing across B2 RNAs (Kolmogorov–Smirnov test values). The X axis represents position (in bases) with regards to the B2 RNA start position for all read k-mers containing an adenosine. (C) Nanopolish signal-density plots at representative B2 positions previously identified as significantly edited (Fig.3D), illustrating group-specific raw-signal differences underlying Xpore estimates of panel A. Nanopolish has been used to aggregate raw nanopore sequencing data into a kmer-level format, which was then used as input to Xpore (see panel A) to estimate the modification rate for a comparison within a single genomic-site and kmer combination. The nanopolish plots displayed here visualize the raw signal distributions that inform the Xpore statistical model of panel A. Any observed differences between the Scamble and AntiAdar distributions correspond to differences in the proportions of non standard bases (i.e. modified bases such as Inosines) between the different samples tested while identical distributions ( as in position 70) would correspond to non-detectable changes in RNA modifications in that position.

These results further support the ability of our customized Illumina-based analysis pipeline in identifying modified Adenosine sites within SINE B2 RNAs in our context.

### B2 RNA editing is increased in hippocampal cells in response to amyloid beta toxicity

In our previous study we had employed a cellular model of amyloid beta toxicity that included inoculation of a hippocampal cell line (HT-22 cells) with amyloid beta peptides (1-42 peptides) (42) and control peptides (reverse 42-1) (R) for 6h (Fig.5A) to test amyloid beta peptide immediate toxicity effects. Thus, we tested whether inoculation of HT22 cells with these peptides could have effects on A-to-I editing levels of B2 RNAs similar to those observed *in vivo*.

**Figure 5.**
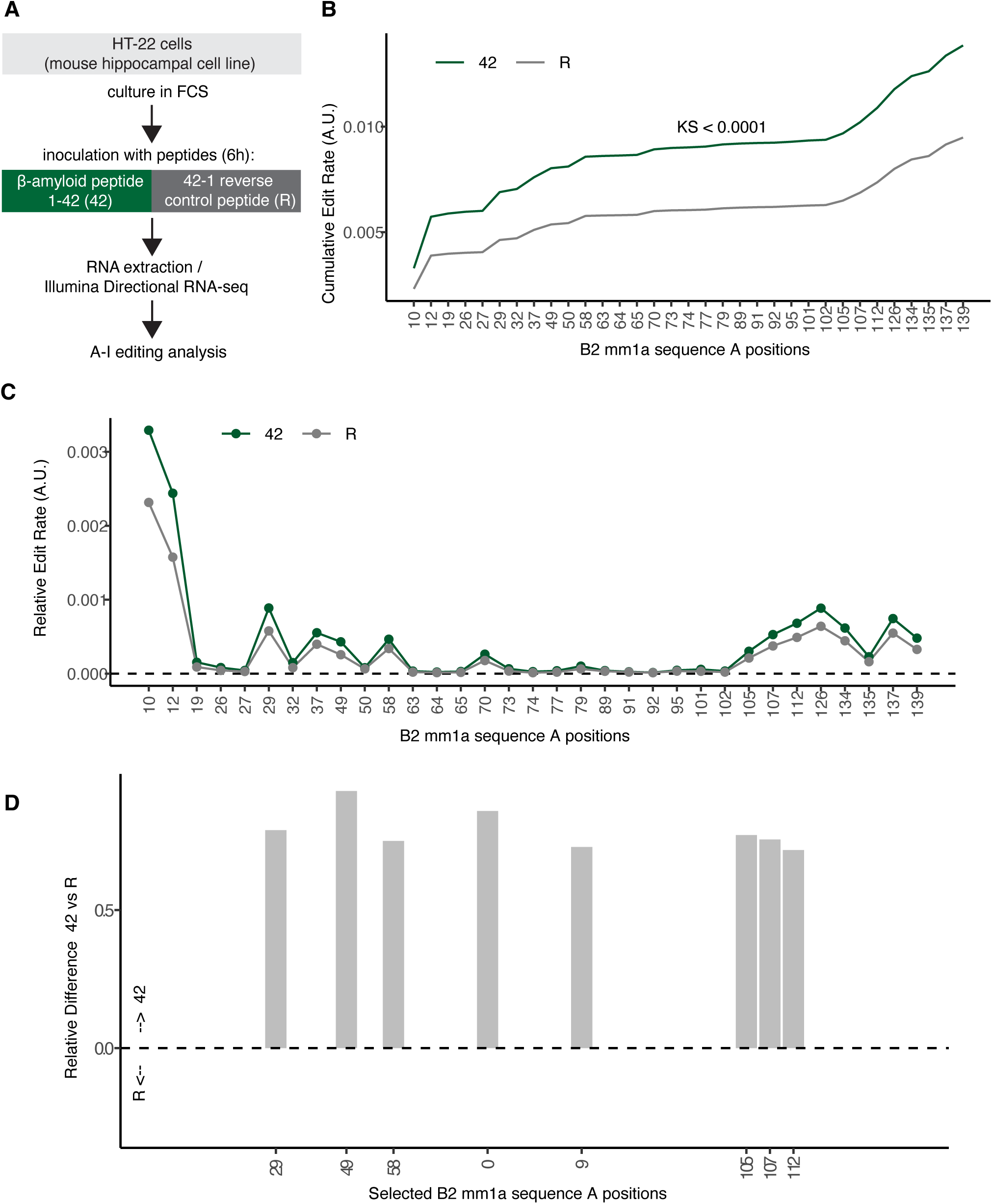
Acute amyloid-beta exposure increases A-to-I editing of B2 RNAs in hippocampal cells. **(A)** Experimental outline of HT-22 cells treated for 6 h with amyloid-β₄₂ peptides (42) or control reverse peptides (R) (n=4) from transcriptome data from (Cheng, Saville, Gollen, Isaac, Mehla, et al., 2020). **(B)** Cumulative distribution of edit rate across B2 RNAs (mm1a) (Kolmogorov–Smirnov test value) (A.U.: arbitrary units). **(C)** Position-specific mean relative editi rates across all consensus A positions (A.U.: arbitrary units). **(D)** Relative differences in edit rate (42 – R) at each position; positive values denote increased editing under amyloid-beta toxicity.

As shown in the edit rate cumulative distributions of Figure 5B, amyloid beta peptides (42) induced a significant increase in total A-to-I editing across B2 RNAs compared to control peptide–treated cells (R) (KS < 0.0001). To determine whether this change reflected specific editing hotspots rather than a global shift, we calculated the relative editing rate at each B2 RNA position.

The resulting position-specific plots (Fig. 5C and D, Suppl. Fig.6) revealed that the editing increase was in alignment with that observed at the same loci identified in AntiAdar and APP^NL-G-F^ vs. WT analyses. In particular, the direction and statistical significance of these changes (Suppl. Fig. 6) mirrored those observed in hippocampal tissue from 3 m and 6 m APP mice (Suppl. Fig. 3), further supporting the consistency of increased edit rate across *in vivo* and *in vitro* models.

**Figure 6.**
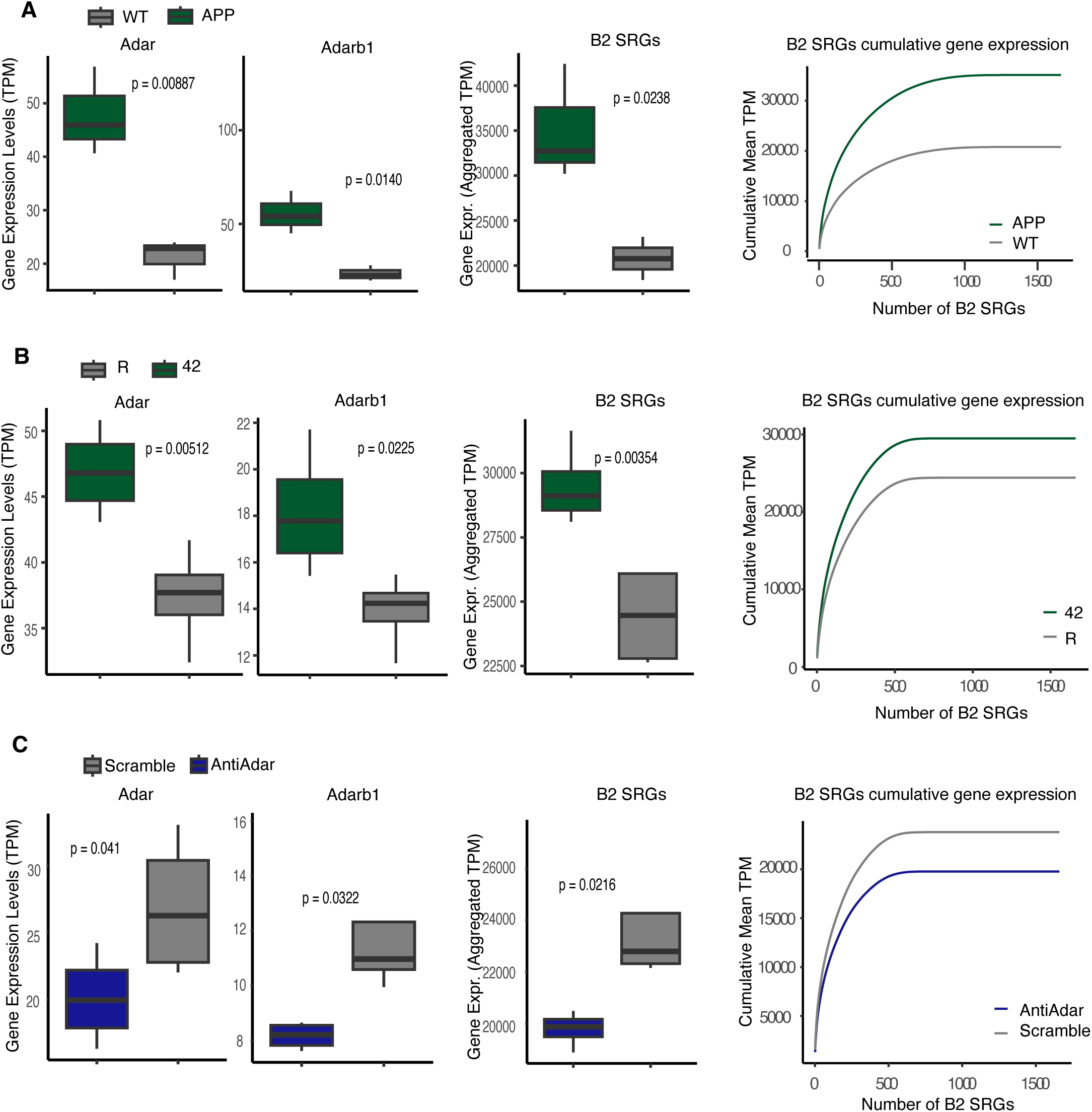
*Adar/Adarb1* expression and B2 stress-response gene levels parallel editing dynamics across models. The first two boxplots in each panel show the expression levels of *Adar* and *Adarb1* The third boxplot shows the aggregate expression (in TPM) of B2 SRGs (1684 genes). The cumulative expression (mean TPM) plot demonstrates that B2 SRG expression follows a continuous distribution not dominated by single transcripts. Expression is in TPM, aggregate TPM, or cumulative TPM as indicated (*P* values, directional Student’s t-test) (A) Expression levels in 6-month-old APP versus WT hippocampi (n=3) of Figure 3. (B) Expression levels in HT-22 cells treated with amyloid beta peptides (42) or a control peptide (R) (n=4) of Figure 5. (C) Expression levels in AntiAdar versus Scramble treated HT-22 cells (n=4) of Figure 2.

Taken together, these results indicate that acute amyloid beta exposure is sufficient to trigger an increase in A-to-I editing of B2 RNAs, recapitulating the editing pattern observed during the early stages of amyloid β pathology in vivo.

### *Adar/Adarb1* levels are increased in response to amyloid beta toxicity and pathology in hippocampal cells

We next examined whether the changes observed in B2 RNAs A-to-I editing were associated with similar alterations in *Adar/Adarb1* expression levels. In hippocampal tissues from APP mice, both *Adar and Adarb1* expression levels were found to be elevated, and this increase was accompanied by higher B2 SRG levels (Fig. 6A). We then asked whether similar relationships could be observed *in vitro*. Indeed, in HT-22 hippocampal cells exposed to amyloid beta toxicity, both *Adar* and *Adarb1* transcript levels were also significantly increased, consistent with the elevated A-to-I editing levels detected in these cells, while the expression levels of B2 SINE RNA-regulated genes (B2 SRGs) were also found to be increased under amyloid beta toxicity conditions (Fig.6B). These changes aligned with changes in B2 RNA editing levels (Suppl. Fig. 7A).

The above findings raised the question whether reducing *Adar/Adarb1* expression could have a suppressive effect on B2 SRG levels, and to investigate this we checked B2 SRG levels during siRNA-mediated *Adar/Adarb1* knockdown in HT-22 cells (AntiAdar) and the subsequent reduction in A to I editing levels (Suppl. Figure 1). As shown in Figure 6C, B2 SRGs transcript levels were significantly lower following *AntiAdar* treatment than in scramble controls, confirming that decreased ADAR activity suppresses B2 SRGs upregulation. These changes also aligned with changes in B2 RNA editing levels (Suppl. Fig. 7B).

Together, these results establish a clear connection between *Adar/Adarb1* expression, amyloid beta toxicity, and B2 SRG activation, suggesting that the ADAR-A-to-I editing machinery may play a regulatory role in B2 RNA-mediated regulation of gene expression in hippocampal cells during amyloid beta toxicity.

## DISCUSSION

Long considered as products of “junk” DNA and transcriptional noise, SINE RNAs have been recently implicated in the cellular response to stress and at the same time designated as primary targets of the A-to-I editing machinery in the cell. This is the first study in hippocampus to identify specific positions within SINE RNAs that act as hotspots of A-to-I editing rather than showing only overall bulk changes across their length. This became possible through our novel analysis strategy, which assesses A-to-I editing of SINE RNAs by approaching these transcripts as a common population of RNA sequences produced by different genomic elements (mapping to the REPome) rather than as a set of separate genomic loci (mapping to the genome). Using this approach we were able to describe distinct position-specific changes in the SINE RNA A-to-I epi-transcriptome in response to amyloid beta pathology and toxicity in hippocampal cells and reveal that editing at specific B2 RNA positions correlates with changes in expression of B2 stress response genes (B2 SRGs). Given the recently identified role of B2 RNA destabilization in the regulation of gene expression and response to cellular stress, these findings unveil a novel link among A-to-I editing, ADAR1 and ADAR2 enzymes, and B2 RNA–regulated gene expression during amyloid beta pathology.

Interestingly, our results demonstrate that editing of SINE RNAs occurs at discrete positions that are reproducible across independent datasets and experimental systems. Instead, we had expected a broadly distributed editing pattern across the entire length of SINE B2 RNAs based on a previously proposed SINE RNA editing mechanism that involves the formation of double stranded RNA molecules from inverted SINE transcripts (Eisenberg & Levanon, 2018; Mehdipour et al., 2020). This new finding raises the potential that intramolecular double strand RNA formations within the SINE B2 RNAs may also play a role in the editing process beyond the mechanism of inverted SINE transcripts.

Using our REPome-based approach, we detected widespread deregulation of B2 RNA editing in amyloid beta pathology, which would have remained undetected by standard genome-based methods. This deregulation was observed both *in vivo*, in mouse hippocampi and brains affected by amyloid beta pathology, and *in vitro*, in hippocampal cells exposed to amyloid beta toxicity. Importantly, the early increase in B2 RNA editing coincided with elevated *Adar/Adarb1* expression levels, suggesting that the ADAR-mediated editing machinery is part of an early epitranscriptomic response to amyloid beta toxicity.

In this study, we focused on the mm1a B2 RNA subfamily, as mm1a—and the closely related mm1t family—most closely match the canonical B2 RNA consensus used in previous B2 RNA functional studies. This ensures that positional editing measurements remain biologically comparable and meaningful, since mm1a elements retain the conserved motifs required for both B2 ribozyme activity and RNA Pol II suppression. Importantly, however, different B2 subfamilies show distinct expression and editing patterns. As illustrated in Supplementary Figure 8, in the case of amyloid beta pathology in mouse models tested in this study, while mm1t editing profiles align with those of mm1a, mm2 elements—harboring several insertions and deletions compared to the previous two subfamilies—display a divergent pattern. These observations indicate that subfamily-specific sequence architectures very likely shape editing behavior, highlighting the need for future functional studies to explore further subfamily-resolved SINE RNA editing.

Our findings raise several new questions. One key issue is whether A-to-I editing serves a protective role—stabilizing B2 RNAs and mitigating their overprocessing—or whether it facilitates B2 RNA turnover and contributes to stress-induced transcriptome remodeling. Distinguishing among these possibilities will require direct experimental modulation of ADAR activity and editing levels in cellular stress-response systems. The coordinated upregulation of *Adar/Adarb1* that we observed in amyloid beta–exposed hippocampal cells suggest a concerted response of the editing machinery, potentially aimed at modulating the stability and regulatory potential of B2 RNAs under cellular stress. Future studies should also examine whether ADAR1 and ADAR2 act redundantly or perform distinct editing functions on B2 RNAs, given their differential subcellular localization and substrate preferences. ADAR2’s role in protein recoding of genes like Gria2 is well accepted, while ADAR1 more prominently suppresses innate immunity, however at the same time, both enzymes demonstrably can edit B2 RNAs, though with potentially different efficiencies (Chalk, Taylor, Heraud-Farlow, & Walkley, 2019). Another important question concerns the mechanistic interplay between RNA editing and the intrinsic ribozyme activity of B2 RNAs. Editing events occurring near or within self-cleavage sites could directly affect RNA stability by altering secondary structures critical for B2 RNA processing. Understanding how editing influences these structural and functional properties will provide insight into how SINE RNAs contribute to cellular resilience or vulnerability during neurodegeneration. Finally, while Nanopore direct RNA sequencing provided an independent support of editing changes detected by short-read sequencing, the type of Nanopore flowcell used cannot unambiguously distinguish inosine from other RNA modifications. Therefore, it cannot be excluded that part of the detected signal variation reflects additional, non-inosine base modifications or potential crosstalk between differing base moieties. For instance, in mRNAs a growing body of evidence suggests that m6A sites and A-to-I editing events are negatively correlated (Xiang et al., 2018) and although B2 RNAs are not known to host m6A moieties, other potential modifications may be impacted by a general loss of ADAR activity.

Overall, elucidating factors, such as ADAR activity, that are upstream of SINE RNA A-to-I editing may provide additional insights regarding the mechanisms underlying the role of SINE RNAs in cell function and related molecular pathologies such as neurodegeneration. This would advance further our understanding of SINE RNAs as potential novel therapeutic targets and not just transcriptional noise. It is notable that Adarb1 (ADAR2) and Adarb2 (ADAR3) genes are in fact B2 RNA-regulated stress response genes themselves (Cheng, Saville, Gollen, Isaac, Belay, et al., 2020), while Adarb1 transcription is amyloid-responsive (Cheng, Saville, Gollen, Isaac, Belay, et al., 2020). By highlighting the site-specific A-to-I editing events in B2 RNAs, our study uncovers the initial results towards a more complete understanding of the complex interacting regulatory axis of A-to-I editing and B2 RNA regulation of stress and amyloid beta toxicity. The human B2 RNA ortholog, SINE Alu RNA (entirely different sequence and structure, but seemingly hosts similar cellular interactions) is also potentially differentially edited in Alzheimer’s disease patients’ hippocampal vasculature (Crooke, Tossberg, Heinrich, Porter, & Aune, 2022). Relatedly, we previously found that SINE Alu RNAs are increasingly processed in Alzheimer’s disease prefrontal cortex and hippocampus samples, which is likely linked to widespread changes in gene expression, relative to age-matched healthy brain tissue (Cheng et al., 2021). Therefore, how the overarching regulatory axis of ADAR regulation and SINE RNA biochemistry interact will also be of key importance to furthering our understanding of neurodegenerative disease such as Alzheimer’s disease.

## MATERIALS AND METHODS

### Cell culture and molecular assays

HT-22 cells were cultured as described previously (Cheng et al., 2020). Transfections of siRNAs against *Adar* and *Adarb1* siRNAs (IDT, see below) were performed through the generation of an equimolar siRNA pool at a10nM final concentration as described before (Cheng et al., 2020) with transfections for 36 hours before harvesting for RNA extraction. For the endonuclease V assays, 250ng total RNA was incubated with 5µL 10X NEB 4 buffer, 1µL endonuclease V enzyme (NEB, M0305S) in a 50µL reaction for 1 hour at room temperature. Untreated samples were topped up with 1µL of nuclease-free water. Quantitative PCR was performed as described before (Cheng et al., 2020) using the same B2 RNA primers. Adar primers used were Forward: 5’ – GCCAAAGACAGTGGTCAACCAG – 3’; Reverse: 5’ – GAACAAGGATGTTGCTGAGGAGC – 3’. Adarb1 primers used were Forward: 5’ – GTATGACGCCAGACTCTCACCA – 3’; Reverse: 5’ - CAGGTCTGGATGCTGGCATTTG - 3’. For normalization of qPCR results, in case of B2 RNA analysis, we used mouse 7sk RNA with primers Forward 5’-GACATCTGTCACCCCATTGA-3’, Reverse 5’-GCCTCATTTGGATGTGTCTG-3’. In case of Adar and Adarb1 we used Hprt with primers Forward 5’-TCCTCCTCAGACCGCTTTT-3’ and Reverse 5’-CCTGGTTCATCATCGCTAATC-3’. IDT siRNAs used in this study are as follows: mm.Ri.Adarb1.13.1 or DsiRNA 5’-AAUCUGAGUCUUUCAGCAUUCUGAC-3 3’-ACUUAGACUCAGAAAGUCGUAAAGACUG-5’, mm.Ri.Adar13.1 or DsiRNA 5’-UCAGUGUUUAUGAUUCCAAAAGACA-3’ 3’-UCAGUCACAAAUACUAAGGUUUUCUGU-5’.

### Nanopore Direct RNA-seq and analysis

For the nanopore data used in Figure 4, RNA from the AntiAdar experiments was pooled from n=4 replicate sets prior to sequencing for each sample type. In the case of mouse hippocampal RNA, 6-month old APP and WT mouse samples were sequenced individually for each replicate. The libraries were prepared according to the vendor-recommended protocol for the SQK_RNA002 direct RNA sequencing kit using 1ug of starting material. Briefly, 9.5uL of RNA was mixed with 1uL RTA adaptor (Nanopore), 1.5uL T4 DNA ligase (NEB, M0202M) and 3uL NEBNext quick ligation buffer (NEB, B6058S), and incubated for 15min at room temperature. The first-strand synthesis reaction was constructed from the ligation product by adding 9uL H2O, 2uL dNTP mix (NEB, N0447S), 8uL FS buffer (Invitrogen, 18080093), 4uL 0.1M DTT (Invitrogen, 18080093) and 2uL Superscript III reverse transcriptase (Invitrogen, 18080093) and incubated at 50C for 50min, 70C for 10min, then placed immediately on ice. The sample was then subjected to a 1.8X bead clean up using the vendor recommended protocol with Mag-Bind TotalPure NGS beads (Omega Bio-Tek, M1378), eluting in 23uL H2O. 6uL of the RMX adaptor (Nanopore), 8uL of the NEBNext quick ligation buffer and 3uL of T4 DNA ligase were added to the eluted RNA:DNA hybrid and incubated for another 15min at room temperature, followed by a 1X bead cleaning reaction. The prepared libraries were loaded and sequenced on a R9.4.1 PromethION flow cell using a PromethION 24 Beta instrument as per manufacturer suggestions.

Nanopore sequencing data was basecalled using Guppy version 3.2.10. Samples were pooled group-wise before mapping. Reads were mapped using minimap2 version 2.24 with the following parameters: -ax sr -uf --secondary=yes. Resulting bam files were then sorted and indexed using samtools-1.2.1. Eventalign tables were generated using Nanopolish 0.14.0 (Loman, Quick, & Simpson, 2015) and modrates were measured using Xpore 2.0 (Pratanwanich et al., 2021) where Nanopolish is a prerequisite for Xpore. Xpore arranges raw signal data from nanopore sequencing by matching sequence kmer and mapping position. We filter this data by removing modification-rate values greater than 0.99 (assumed to be an artifact of Xpore’s model since 100% editing rate should not occur biologically), we also filter for kmer positions containing an A, to give a suitable substrate for A-to-I editing, finally we filter to remove absolute value of differential modification rate less than 0.2 to ensure only strongly modified sites remain. This data is used to produce the Xpore signal plots in Figure 4A and 4B. To produce raw signal plots (Figure 4C), data was aligned using minimap2 with the same version and parameters on ensemble release 102 Mus musculus genome (mm10). Samtools was used as previously described to sort and index the bam files. Nanopolish 2.1 was run on these bam files. Then the coordinate system from these nanopolish tables were intersected with the coordinates of B2 Mm1a in the genome using bedtools v2.31.1 with the -wo flag (Quinlan & Hall, 2010). The resulting nanopolish table, now only containing nanopolish signal which intersects B2 loci, was used to generate raw signal plots in Figure 4C.

### Bioinformatics analysis SINE RNA A to I editing

A custom reference was constructed from unique B2 elements in the mouse genome, hence called the REPome (unique B2ome). For the construction of the “REPome” we retrieved B2 sequences for all identified B2 repeats (mm1a, mm1t and mm2) within the range of 125 bp or longer in order to exclude truncated forms from UCSC mm10 Repeat Masker (August 2018) and subsequently collapsed sequences present in our list more than once into one unique representative sequence, resulting only in unique B2 sequences. RNA-seq reads for each sample were mapped to the REPome using bwa-0.7.19 (H. Li & Durbin, 2009) with the following parameters: aln, with maximum ten (10) allowed mismatches. The goal of aligning to the unique REPome was to ensure that each element is present only once in the reference. After alignment, files were converted to bam format, sorted and indexed using samtools-1.21 (H. Li et al., 2009). Then, the sense-aligning fraction was extracted from the samfile (samtools flag -F 16). Subsequently, the RNA edit detection software REDItools v1.2.1 (denovo script) (Picardi & Pesole, 2013) was utilized to produce tables of base alignments for each file. This software was run using the following modified parameters: -c 1, -q 1, -m 1, -O 0, -V 1, -n 0. The rational for the selection of these parameters was as follows; -c denotes the minimum read coverage, which was set to 1 as alignment on the unique reference may produce sparse read-coverage; -q denotes minimum quality score, which was set to 1 as highly edited RNA may appear to the sequencer as low quality; -m is mapping quality score with the same rationale as previous; -O is min homopolymer length, which was set to zero so that REDItools will treat homopolymeric regions of the read identically to the heterogeneous regions; -n is the minimum editing frequency, set to 0 for maximum sensitivity.

After the REDItools tables were produced, we merged them with file identifiers into one single table. Then we computed the editing rate as follows. Initially, we constructed two sets of proportion statistics from base frequences. Within each sample and for each position (1-180) of the B2 RNAs, we computed the counts of all reference A, T, C, and G respectively, terming these “expected” counts. We repeated this count for all bases that are present in the reads, terming these “observed” counts. These counts are normalized to the sum of all expected and observed bases, respectively, yielding proportions. Thus, for each sample and B2 position, we generated two sets of proportion statistics, each of which was composed of four proportions A, T, C, G (sum to 1). The first set was the “expected” proportions, which were generated from the expressed (>0 read coverage) regions of the unique B2-ome, without weighting the reference base contribution by the number of reads. The second set was the “observed” proportions, which were generated from the bulk aligned reads regardless of which sequence they aligned to. These proportions were used in the calculation of the editing rate per sample and position, which was (obsGprop/(obsGprop+obsAprop))/expA. This editing rate was based on the canonical G/(A+G), however it utilized the population-level observed proportions previously described. Lastly, the editing rate was normalized to the number of unique reference regions which have an A at the given position. Additionally, we calculated this rate separately for each B2 subfamily, filtered for only B2 Mm1a elements, and then we examined only positions within the B2 Mm1a consensus sequence that contain an A.

The editing rate in Supplementary Figure 1C was calculated using the raw reditools table. We filtered this table for positions which have an A in the reference, then computed the editing rate as G/(A+G) where these letters refer to the counts of reads aligning to that position. Then, we computed the mean of this edit rate across positions and samples to obtain a per-sample per-position rate in parity with our method.

### Gene expression analysis

B2 stress response genes are genomic loci that have been shown to be regulated by B2 RNA. The list of gene names was retrieved from our previous study (Cheng et al., 2020). To calculate the gene expression levels we first aligned fastq files against Ensembl release 102 Mus Musculus genome GRCm38 using Hisat2 v2.2.0 with the –dta option. The gene expression levels were calculated by using Stringtie 2.2.1 with -G -A -B -e parameters in conjunction with transcriptome GTFs of the same Ensembl release. TPM levels from the analyzed gene or set of genes were extracted for each sample from the resulting abundance files.

### Transcriptome data sources and data access

Raw Illumina RNA sequencing data (fastq files) for the APP/WT mouse and HT-22 transcriptome data originate from our previous study (Cheng, Saville, Gollen, Isaac, Mehla, et al., 2020) and are deposited in GEO with access number GSE149243. Immunohistochemistry and behavioral studies of this mice model are described in our previous study (Mehla et al., 2019). The tested mice carry the Arctic, Swedish, and Beyreuther/Iberian mutations (APPNL-G-F/NL-G-F) and were provided by the RIKEN Center for Brain Science, Japan; the colony was maintained at the University of Lethbridge vivarium. The external mouse dataset comprises RNA sequencing data from mice with human transgenes for mutant forms of APP and PSEN1, as well as wild type controls. The dataset was retrieved from the GEO access number GSE136861 series.

Novel Illumina and Nanopore RNA data of this study has been deposited to GEO with access number GSE316378.

## ACKNOWLEDGEMENTS

This work has been supported by the #PJT180336 grant from CIHR to AZ and MM, a Grant # 201900003 to AZ and MM from the Alzheimer Society of Alberta and Northwest Territories and the Alberta Prion Research Institute, a Discovery Grant # RGPIN-2018-05955 to AZ from NSERC, the Levmax 222301783 grant from Alberta innovates, Alzheimer Association AARG Grant #23-1152151, CIHR Project Grant # 507080 to MHM, and a Compute Canada Resource Allocation Grant to AZ. AZ has been supported by the Canada Research Chairs Program and is a former EMBO and DFG long-term fellow. YC, LS and LM have been supported by an Alberta Innovates graduate fellowship. RR and TH have been supported by the NSERC CREATE RNA Innovation program. We are grateful to the RFHS Statistical Genomics Platform and the CCMB Bioinformatics Core for their help. We are grateful to Dr. Angeliki Pantazi for extensively reviewing, editing and commenting on the manuscript.

## AUTHOR CONTRIBUTIONS

LM: Bioinformatics analysis, statistical analysis, establishment and testing of the analysis pipelines, data visualization and writing of the manuscript; LS: testing of the next-generation-sequencing and in vitro assay pipelines, library construction; sequencing; TH, RR, CT: in vitro assays and analysis; YC: Bioinformatics analysis; IK: data interpretation; MM: data interpretation, manuscript edit; AZ: conception and design, establishment and testing of data generation and analysis pipelines, bioinformatics analysis, data interpretation, data visualization and writing of the manuscript, overall supervision.

