## Supplementary for "Differential adenosine to inosine RNA editing of SINE B2 non-coding RNAs in mouse unveils a novel type of epi-transcriptome response to amyloid beta neuro-toxicity"

### SUPPLEMENARY FIGURES (8 FIGURES)

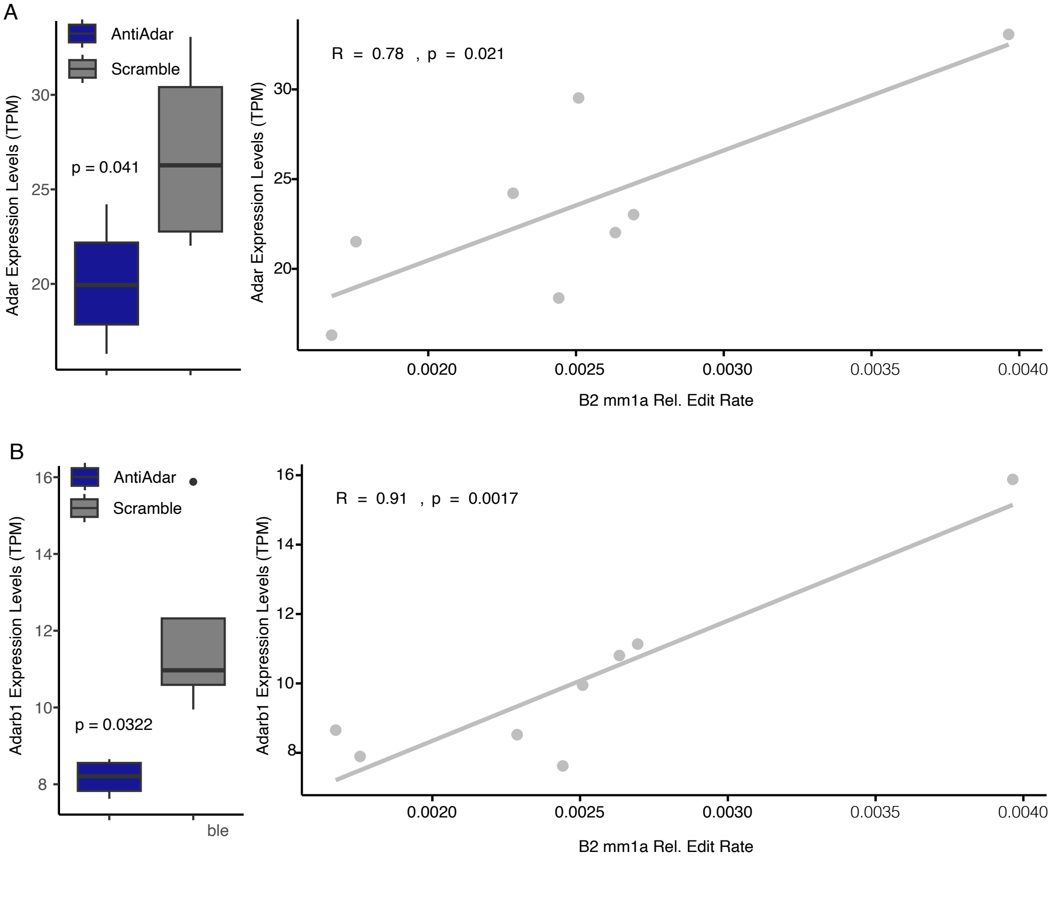


SUPPLEMENTARY FIGURE 1

**Supplementary Figure 1.**

Sample-wise correlations between Adar/Adarb1 expression and A-to-I edit rates in siRNA-treated HT22 cells (see Figure 2). Left panels show transcript per million (TPM) expression of *Adar* (**A**) and *Adarb1* (**B**) as measured through RNA-seq.

*P*-values in (A) are derived from one-tailed Student's *t*-test. Right panels display per replicate Pearson correlations between *Adar* or *Adarb1* transcript abundance and cumulative edit rate across the subset of highly edited (selected) A positions (Panel A: R=0.78, p=0.021; Panel B: R=0.91, p=0.0017).

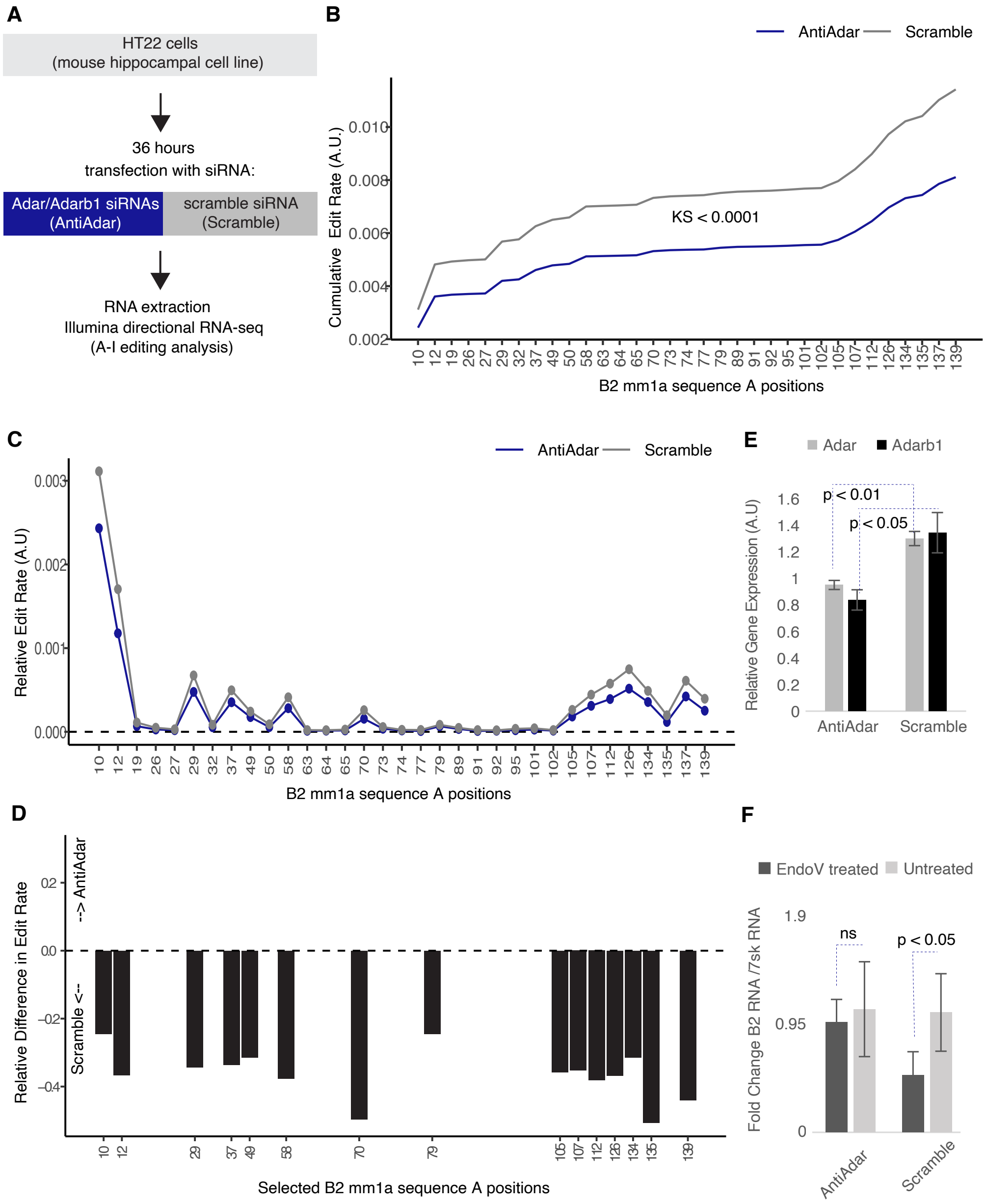
A
HT22 cells
(mouse hippocampal cell line)
36 hours
transfection with siRNA:
Adar/Adarb1 siRNAs
(AntiAdar)
scramble siRNA
(Scramble)
RNA extraction
Illumina directional RNA-seq
(A-I editing analysis)
B
AntiAdar
Scramble
Cumulative Edit Rate (A.U.)
KS < 0.0001
B2 mm1a sequence A positions
C
AntiAdar
Scramble
Relative Edit Rate (A.U)
B2 mm1a sequence A positions
D
Relative Difference in Edit Rate
--> AntiAdar
Scramble <--
Selected B2 mm1a sequence A positions
E
Adar
Adarb1
Relative Gene Expression (A.U)
p < 0.01
p < 0.05
AntiAdar
Scramble
F
EndoV treated
Untreated
Fold Change B2 RNA /7sk RNA
ns
p < 0.05
AntiAdar
Scramble

FIGURE 2

**Supplementary Figure 2.**

Our customized approach for calculating A-to-I edit rates achieves higher fidelity than standard genome-aligned methods. Two methodological distinctions underlie this improvement. In typical Illumina-based A-to-I editing detection, reads are aligned to the genome and intersected with annotated elements of interest (“alignment on genome”). In contrast, our method aligns reads directly to the unique transcriptome of the elements of interest (“alignment on unique B2”; panel A). A second distinction lies in the stage at which values are aggregated. The standard approach calculates editing rate per reference transcript and subsequently aggregates these across transcripts to derive a sample-level value (“edit-then-group”). Our method first aggregates the transcripts and then computes the editing rate across them collectively (“group-then-edit-rate”). Comparisons were performed using the same data as Figure 2 (n=4 per group). Alignment on genome was performed using the same alignment parameters as for unique B2, see methods.

**(A)** Differences in the alignment strategy. Genome-based alignment yields reduced read coverage across repetitive B2 elements, whereas alignment to the unique B2-ome achieves markedly higher base coverage.

**(B)** The customized “group-then-edit-rate” approach, as applied throughout the study (identical to Figure 2B – presented here to facilitate comparison with panel C), reveals reduced editing in AntiAdar compared to Scramble samples, consistent with qPCR results for *Adar/Adarb1* mRNA levels (Figure 2E,F and Nanopore data – Figure 4).

**(C)** The same dataset analyzed using the typical approach. Editing rate is computed using the conventional A-to-I formula *G/(A+G)* across reference positions containing an A. The overall trend diverges from *Adar/Adarb1* expression patterns, showing minimal difference between conditions and even a mild apparent increase in AntiAdar samples, which are expected to exhibit lower editing. There is also no positional divergence in editing rates across the different positions.

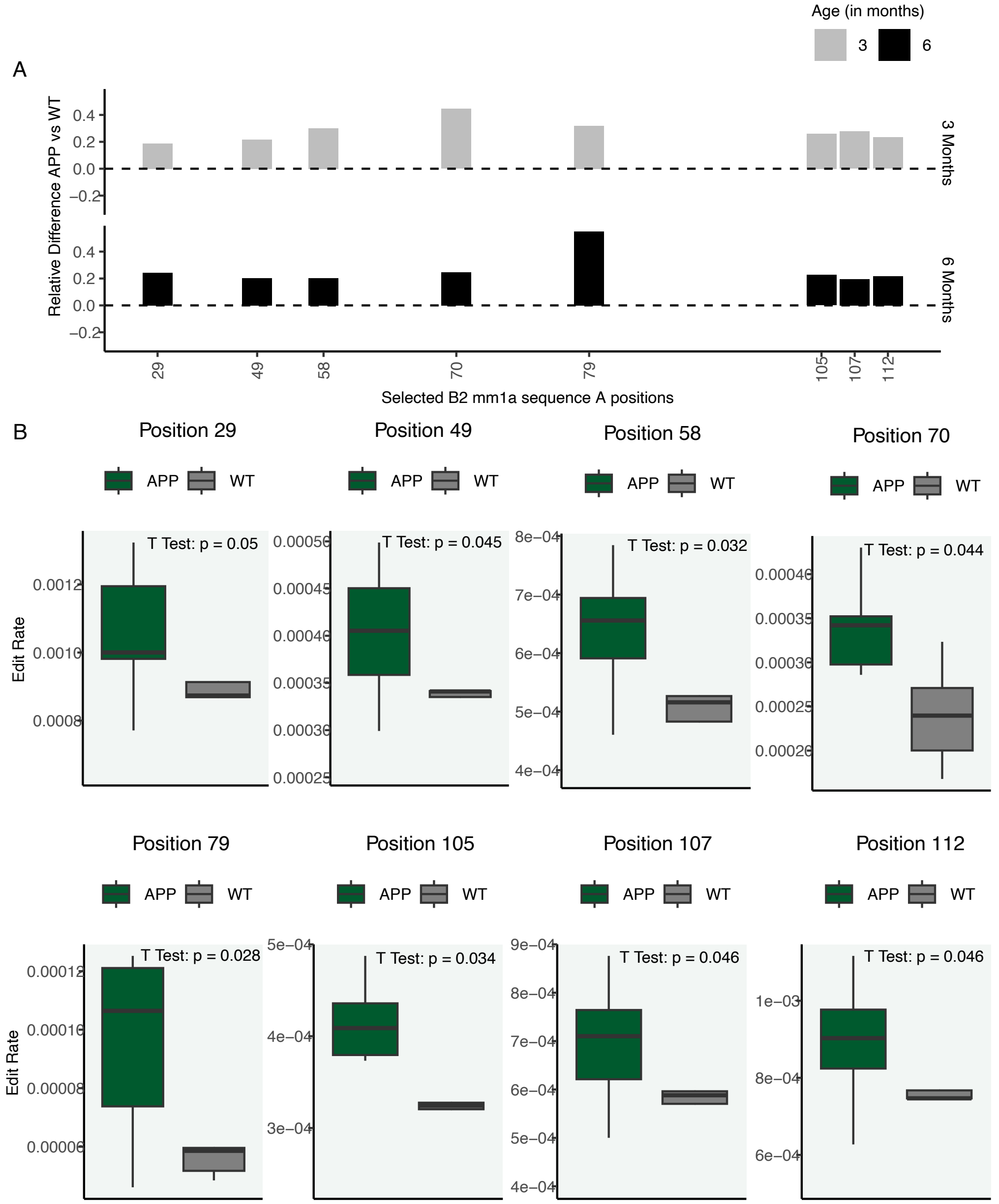


SUPPLEMENTARY FIGURE 3

**Supplementary Figure 3.**

**(A)** Relative differences in edit rate at selected subset positions in 3- and 6-month-old mice (n=3 for each treatment at each age group, except for 3mo WT mice which used n=2). Positions were chosen from the Adar knockdown dataset (Figure 2C) according to two criteria: (i) they lie within the main body of the B2 element (approximately before position 120 nt) to avoid poly-A tail regions that may introduce artefacts, and (ii) they exhibit high editing magnitude in the knockdown data. Although all positions in Figure 2C correspond to adenosines in the consensus Mm1a sequence, editing magnitude varies markedly across sites. The subset shown here represents those positions with a relative edit rate > 0.0001 with a statically significant difference in editing (see panel B below) and is used in all subsequent supplementary analyses (Figure is partially identical with Fig.3 upper two panels and presented here to enable comparison with panel B below).

**(B)** Boxplots comparing the above subset-site editing between APP and WT mice at 3 and 6 months. Both ages show elevated editing under disease conditions, consistent with Figure 3B. *P* values were obtained by a one-tailed Student's t-test. Box plots are produced without outliers.

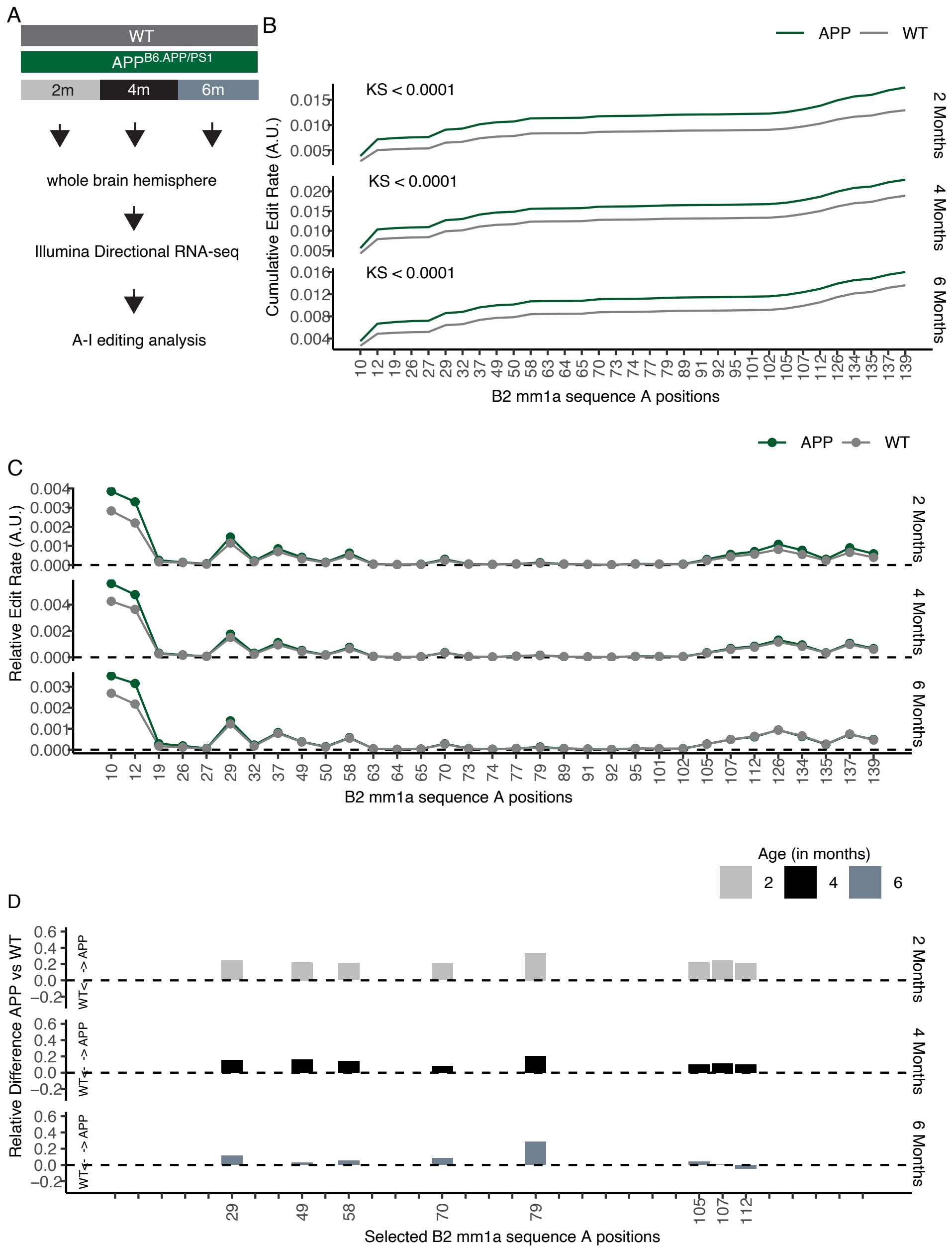


SUPPLEMENTARY FIGURE 4

**Supplementary Figure 4.**

Validation of editing dynamics in an independent mouse model expressing human APP and PSEN1 transgenes (GEO GSE136861). This dataset includes whole-brain RNA-seq from APP and WT mice aged 2 (n= 10, 9 respectively), 4 (n= 5, 9 respectively), and 6 (n= 7, 13 respectively) months to match the primary dataset (Figure 3). The overall pattern parallels our observations in Figure 3: APP mice exhibit increased editing.

**(A)** Schematic overview of the external dataset and processing workflow.
(**B**) Cumulative distribution of edit rate across B2 RNAs (mm1a) per position in APP versus WT groups across the three age ranges. Y-axis indicates mean cumulative editing rate; statistical values are from Kolmogorov–Smirnov tests.
(**C**) Position-specific mean relative edit rates across all consensus A positions (A.U.: arbitrary units).

(**D**) Relative editing differences (APP − WT) along the B2 Mm1a sequence, with positive values denoting higher editing in APP.

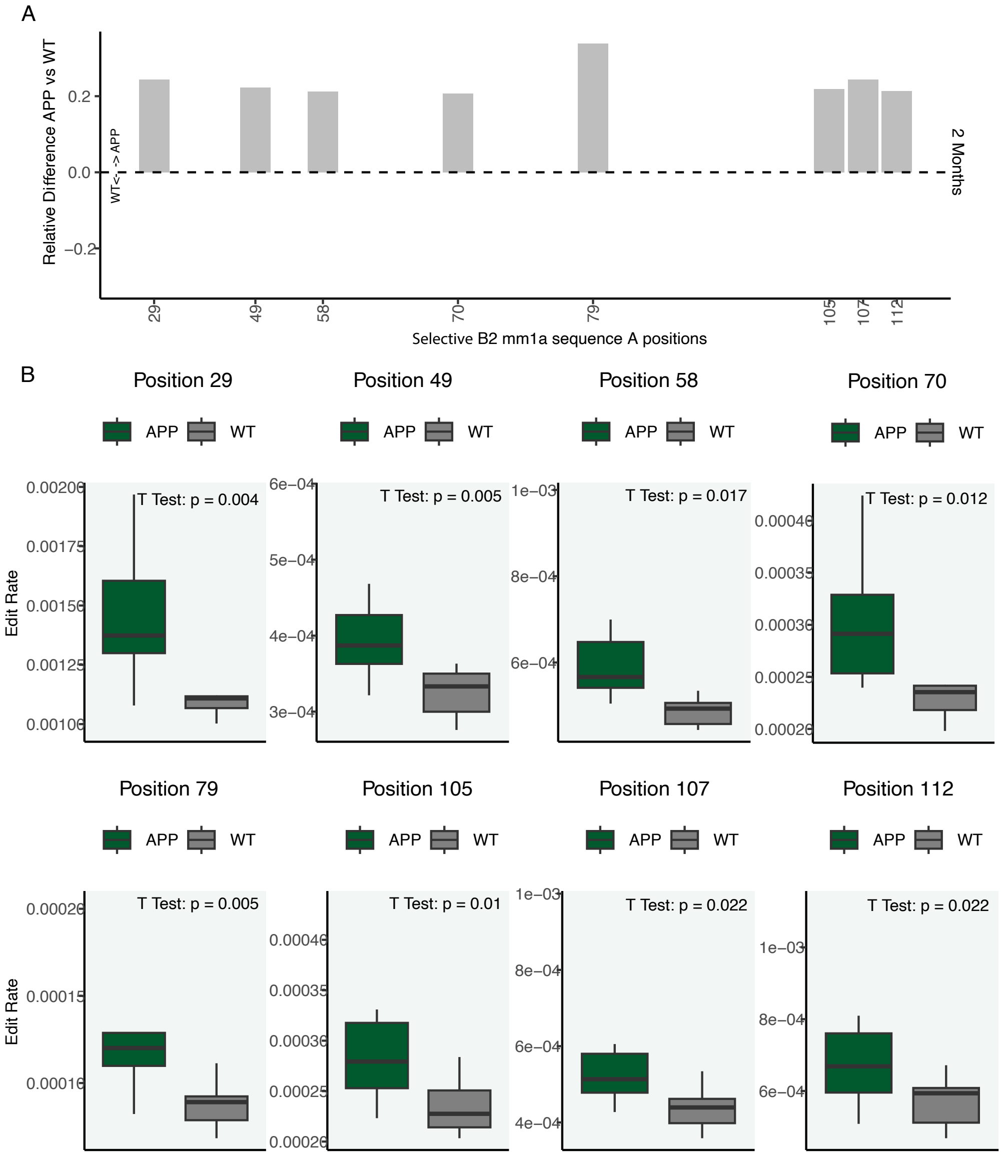


SUPPLEMENTARY FIGURE 5

**Supplementary Figure 5.**

**(A)** Relative editing differences at subset positions (as defined in Supplementary Figure 3) in 2-month-old mice from the external dataset (see Supplementary Figure 4) (n= 10 and 9 for APP and WT, respectively). Figure is partially identical with Suppl. Fig.4 upper panel and presented here to enable comparison with panel B below.

**(B)** Boxplots comparing subset positions between APP and WT groups at 2 months. *P* values were obtained using one-tailed Student's *t*-tests.

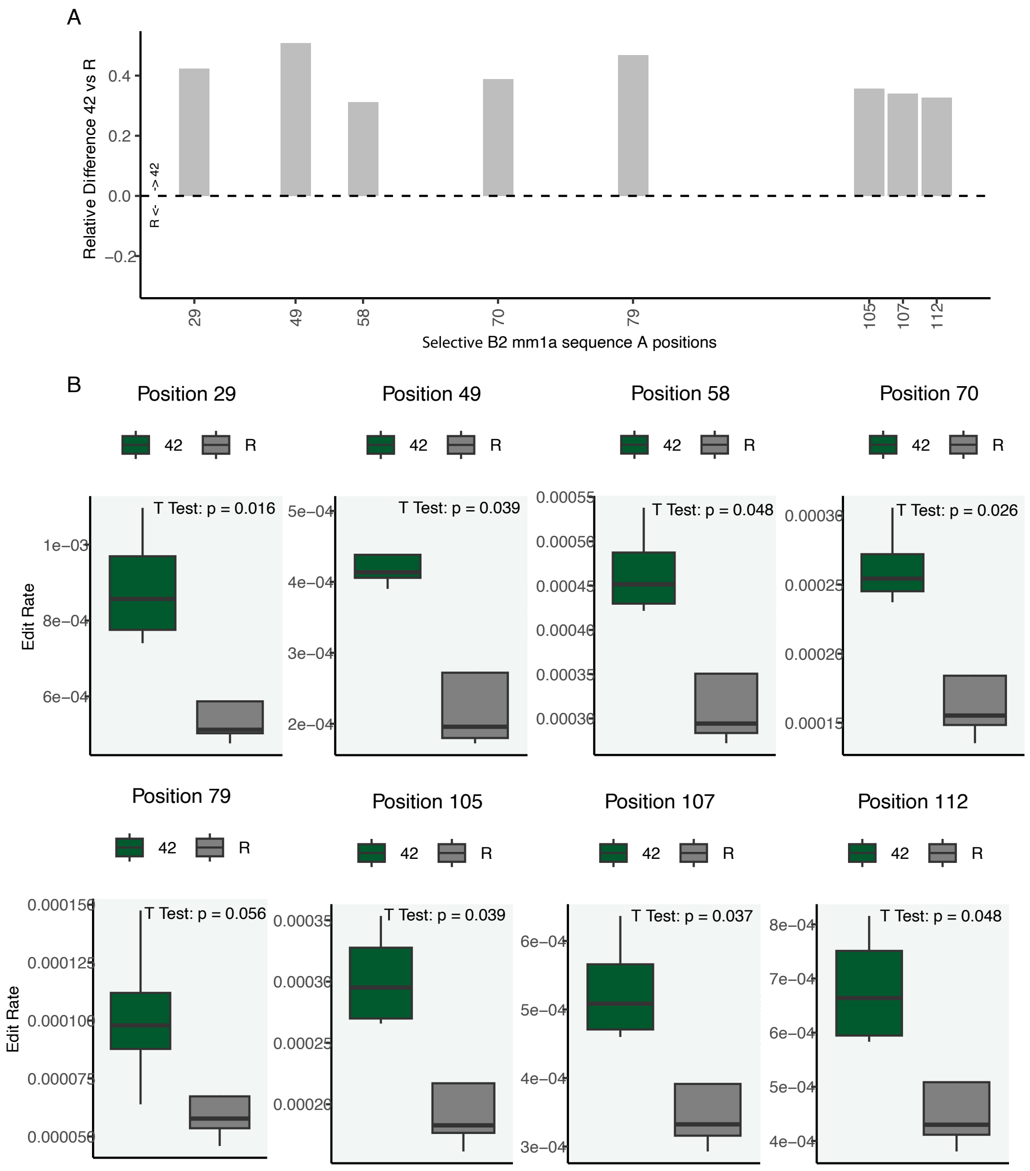


SUPPLEMENTARY FIGURE 6

**Supplementary Figure 6.**

Subset-site editing changes in HT22 cells exposed to amyloid-β toxicity (see Figure 5).

(**A**) Relative editing differences at subset positions. Supplementary Figure 3A legend describes how positions have been selected. (Figure is identical with Fig.5C upper and presented here to enable comparison with panel B below).

(**B**) Boxplots comparing amyloid-treated (42) and reverse-peptide-treated (R) cells. *P* values are from one-tailed Student's *t*-tests.

A

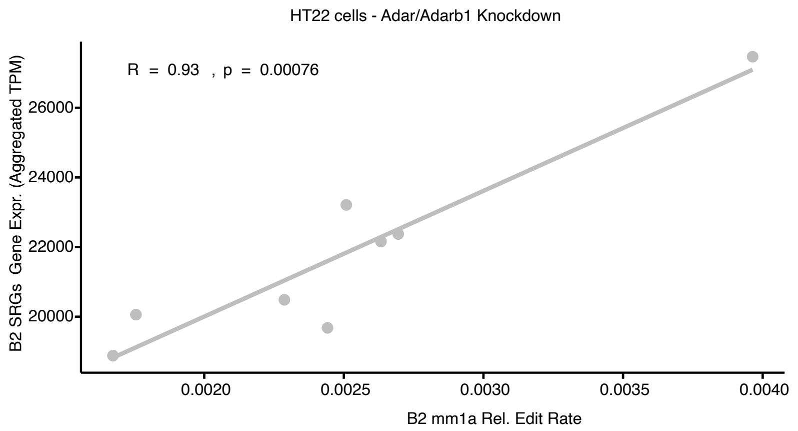


B

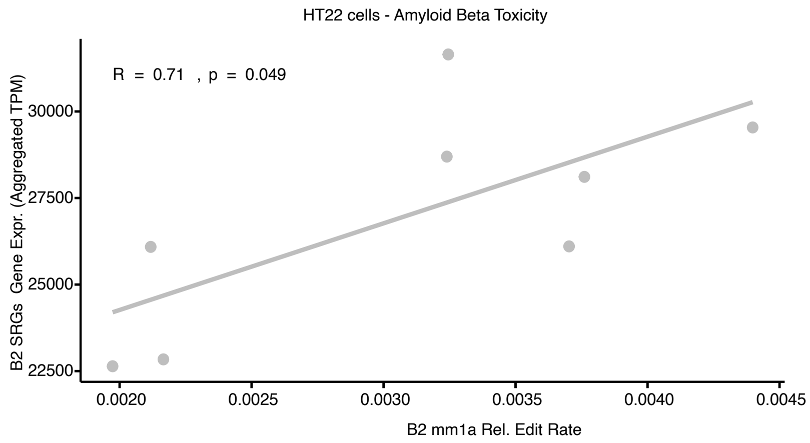


SUPPLEMENTARY FIGURE 7

**Supplementary Figure 7.**

Sample-wise correlation between B2-SRG expression and A-to-I editing in **(A)** siRNA-knockdown (Figure 2 dataset) and **(B)** during response to amyloid beta toxicity (Figure 5 dataset) in HT22 cells, showing per replicate Pearson correlations between cumulative B2-SRG expression and total editing at the subset positions defined in Supplementary Figure 3 (Panel A: $R=0.93, p=0.00076$; Panel B: $R=0.71$, $p=0.0049$). The positive correlations across both experiments support a tight coupling between SINE-derived gene activity and A-to-I editing levels in HT22 cells.

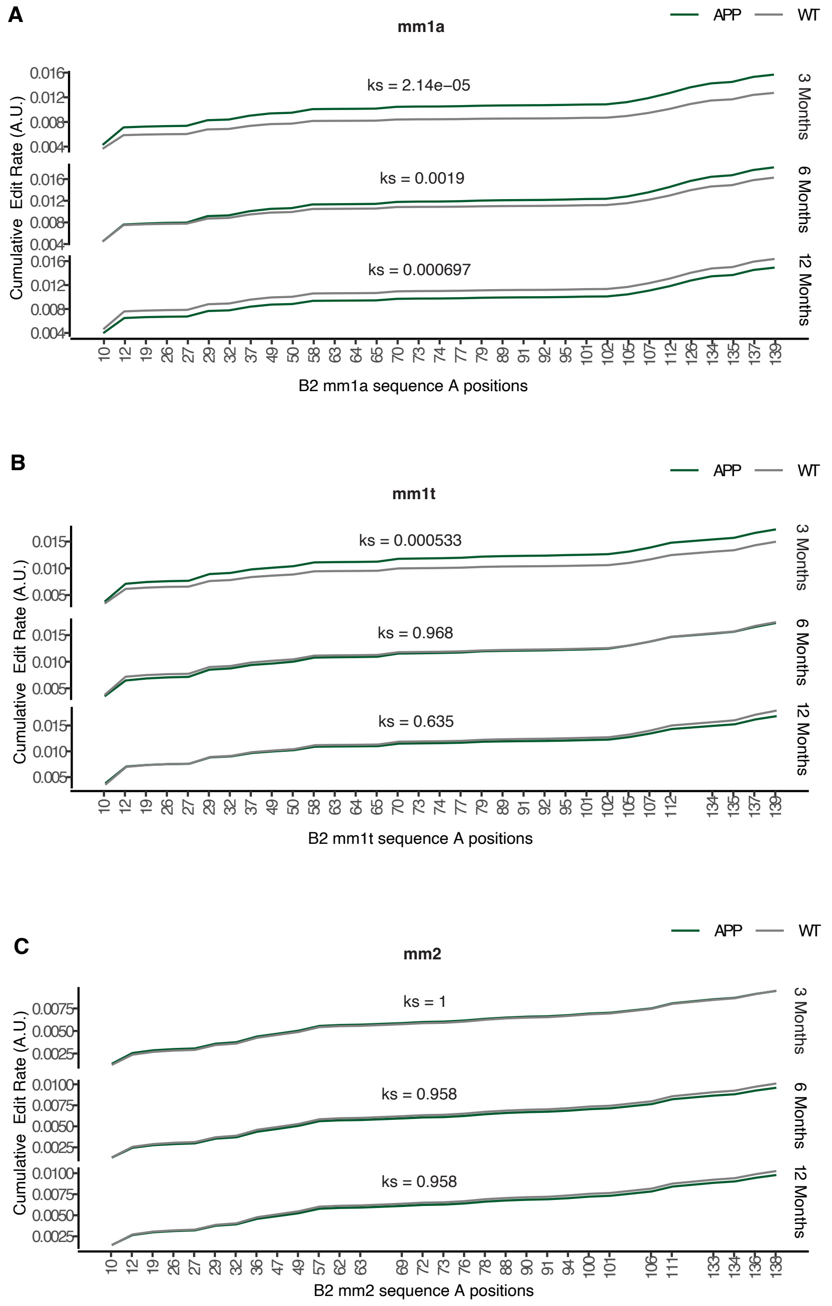


SUPPLEMENTARY FIGURE 8

**Supplementary Figure 8**

Cumulative edit rate between APP and WT samples (as described in Fig.3B) across three different B2 RNA subfamilies.

**(A)** Cumulative edit rate specific to B2 mm1a subfamily.
**(B)** Cumulative edit rate specific to B2 mm1t subfamily.
**(C)** Cumulative edit rate specific to B2 mm2 subfamily.
